# Expanding diversity of Asgard archaea and the elusive ancestry of eukaryotes

**DOI:** 10.1101/2020.10.19.343400

**Authors:** Yang Liu, Kira S. Makarova, Wen-Cong Huang, Yuri I. Wolf, Anastasia Nikolskaya, Xinxu Zhang, Mingwei Cai, Cui-Jing Zhang, Wei Xu, Zhuhua Luo, Lei Cheng, Eugene V. Koonin, Meng Li

**Affiliations:** Shenzhen Key Laboratory of Marine Microbiome Engineering, Institute for Advanced Study, Shenzhen University, Shenzhen, Guangdong, 518060, P. R. China; National Center for Biotechnology Information, National Library of Medicine, National Institutes of Health, Bethesda, Maryland 20894, USA; State Key Laboratory Breeding Base of Marine Genetic Resources, Key Laboratory of Marine Genetic Resources, Fujian Key Laboratory of Marine Genetic Resources, Third Institute of Oceanography, State Oceanic Administration, Xiamen 361005, P. R. China; Key Laboratory of Development and Application of Rural Renewable Energy, Biogas Institute of Ministry of Agriculture, Chengdu 610041, P.R. China

## Abstract

Comparative analysis of 162 (nearly) complete genomes of Asgard archaea, including 75 not reported previously, substantially expands the phylogenetic and metabolic diversity of the Asgard superphylum, with six additional phyla proposed. Phylogenetic analysis does not strongly support origin of eukaryotes from within Asgard, leaning instead towards a three-domain topology, with eukaryotes branching outside archaea. Comprehensive protein domain analysis in the 162 Asgard genomes results in a major expansion of the set of eukaryote signature proteins (ESPs). The Asgard ESPs show variable phyletic distributions and domain architectures, suggestive of dynamic evolution via horizontal gene transfer (HGT), gene loss, gene duplication and domain shuffling. The results appear best compatible with the origin of the conserved core of eukaryote genes from an unknown ancestral lineage deep within or outside the extant archaeal diversity. Such hypothetical ancestors would accumulate components of the mobile archaeal ‘eukaryome’ via extensive HGT, eventually, giving rise to eukaryote-like cells.

## Introduction

The Asgard archaea are a recently discovered archaeal superphylum that is rapidly expanding, thanks to metagenomic sequencing (*1*–*5*). The Asgard genomes encode a diverse repertoire of eukaryotic signature proteins (ESPs) that far exceeds the diversity of ESPs in other archaea. The Asgard ESPs are particularly enriched in proteins involved in membrane trafficking, vesicle formation and transport, cytoskeleton formation and the ubiquitin network, suggesting that these archaea possess a eukaryote-type cytoskeleton and an intracellular membrane system (*2*).

The discovery of the Asgard archaea rekindled the decades old but still unresolved fundamental debate on the evolutionary relationship between eukaryotes and archaea that has shaped around the ‘2-domain (2D) versus 3-domain (3D) tree of life’ theme (*6*–*8*). The central question is whether the eukaryotic nuclear lineage evolved from a common ancestor shared with archaea, as in the 3D tree, or from within the archaea, as in the 2D tree. The discovery and phylogenomic analysis of Asgard archaea yielded strong evidence in support of the 2D tree, in which eukaryotes appeared to share common ancestry with one of the Asgard lineages, Heimdallarchaeota (*1*, *2*, *5*). However, the debate is not over as arguments have been made for the 3D topology, in particular, based on the phylogenetic analysis of RNA polymerases, some of the most highly conserved, universal proteins (*9*, *10*).

Molecular phylogenetic methods alone might be insufficient to resolve the ancient ancestral relationship between archaea and eukarya. To arrive at a compelling solution, supporting biological evidence is crucial (*11*), because, for example, the transition from archaeal, ether-linked membrane lipids to eukaryotic, ester-linked lipids that constitute eukaryotic (and bacterial) membranes (*12*) and the apparent lack of the phagocytosis capacity in Asgard archaea (*13*) are major problems for scenarios of the origin of eukaryotes from Asgard or any other archaeal lineage. Some biochemical evidence indicates that Asgard archaea possess an actin cytoskeleton regulated by accessory proteins, such as profilins and gelsolins, and the endosomal sorting complex required for transport machinery (ESCRT) that can be predicted to function similarly to the eukaryotic counterparts (*14*–*16*). Generally, however, the biology of Asgard archaea remains poorly characterized, in large part, because of their recalcitrance to growth in culture (*17*). To date, only one Asgard archaeon, *Candidatus* Prometheoarchaeum syntrophicum strain MK-D1, has been isolated and grown in culture (*17*). This organism has been reported to form extracellular protrusions that are involved in its interaction with syntrophic bacteria, but no visible organelle-like structure and, apparently, little intracellular complexity.

The only complete, closed genome of an Asgard archaeon also comes from *Candidatus* P. syntrophicum strain MK-D1 (*17*) whereas all other genome sequences were obtained by binning multiple metagenomics contigs. Furthermore, these genome sequences represent but a small fraction of the Asgard diversity that has been revealed by 16S rRNA sequencing (*18*, *19*).An additional challenge to the study of the relationship between archaea and eukaryotes is that identification and analysis of the archaeal ESPs are non-trivial tasks due to the high sequence divergence of many if not most of these proteins. At present, the most efficient, realistic approach to the study of Asgard archaea and their eukaryotic connections involves obtaining high quality genome sequences and analyzing them using the most powerful and robust of the available computational methods.

Here we describe metagenomic mining of the expanding diversity of the superphylum Asgard, including the identification of six additional phylum-level lineages that thrive in a wide variety of ecosystems and are inferred to possess versatile metabolic capacities. We show that these uncultivated Asgard groups carry a broad repertoire of ESPs many of which have not been reported previously. Our in depth phylogenomic analysis of these genomes provides insights into the evolution of Asgard archaea but calls into question the origin of eukaryotes from within Asgard.

## Results

### Reconstruction of Asgard archaeal genomes from metagenomics data

We reconstructed 75 metagenome-assembled genomes (MAGs) from the Asgard superphylum that have not been reported previously. These MAGs were recovered from various water depths of the Yap trench, intertidal mangrove sediments of Mai Po Nature Reserve (Hong Kong, China) and Futian Mangrove Nature Reserve (Shenzhen, China), seagrass sediments of Swan Lake Nature Reserve (Rongcheng, China) and petroleum samples of Shengli oilfield (Shandong, China) (Supplementary Table 1, Supplementary Figure 1). For all analyses described here, these 75 MAGs were combined with 87 publicly available genomes, resulting in a set of 162 Asgard genomes. The 75 genomes reconstructed here were, on average, 82% complete and showed evidence of low contamination of about 3%, on average (Supplementary Figure 2).

### Classification of Asgard genes into clusters of orthologs

The previous analyses of Asgard genomes detected a large fraction of “dark matter” genes (*20*). For example, in the recently published complete genome of *Candidatus* Prometheoarchaeum syntrophicum, 45% of the proteins are annotated as “hypothetical”. We made an effort to improve the annotation of Asgard genomes by investigating this dark matter in greater depth, and developing a dedicated platform for Asgard comparative genomics. To this end, we constructed Asgard Clusters of Orthologous Genes (asCOGs) and used the most sensitive available methods of sequence analysis to annotate additional Asgard proteins, attempting, in particular, to expand the catalogue of Asgard homologs of ESPs (see Materials and Methods for details).

Preliminary clustering by sequence similarity and analysis of the protein cluster representation across the genomes identified the set of 76 most complete Asgard MAGs (46 genomes available previously and 30 ones reported here) that cover most of the group diversity (Supplementary Table 1). The first version of the asCOGs presented here consists of 14,704 orthologous protein families built for this 76-genome set. The asCOGs cover from 72% to 98% (92% on average) of the proteins in these 76 genomes (additional data file 1). Many asCOGs include individual domains of large, multidomain proteins.

The gene commonality plot for the asCOGs shows an abrupt drop at the right end, which reflects a surprising deficit of nearly universal genes (Fig. 1). Such shape of the gene commonality curve appears anomalous compared to other major groups of archaea or bacteria with many sequenced genomes (*21*). For example, in the case of the TACK superphylum of archaea, for which the number of genomes available is similar to that for Asgard, with a comparable level of diversity, the commonality plot shows no drop at the right end, but instead, presents a clear uptick, which corresponds to the core of genes represented in (almost) all genomes (Fig. 1). Apparently, most of the Asgard genomes remain incomplete, such that conserved genes were missed randomly. Currently, there are only three gene families that are present in all Asgard MAGs, namely, a Zn-ribbon domain, a Threonyl-tRNA synthetase and an aminotransferase (additional data file 1).

**Figure 1.**
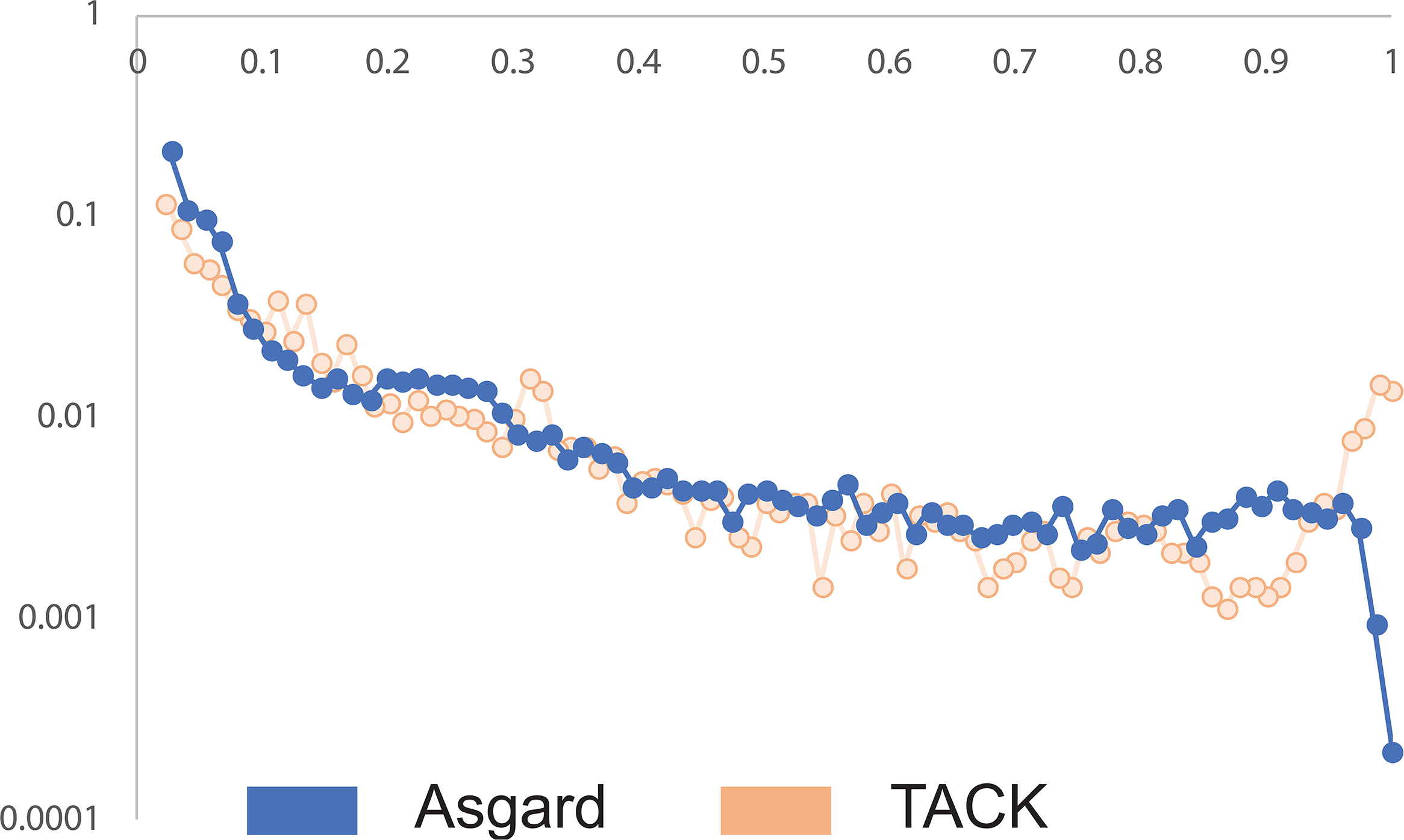
Gene commonality plots for Asgard archaea and the TACK superphylum. The gene commonality plot showing the number of asCOGs in log scale (Y-axis) that include the given fraction of analyzed genomes (X-axis). The Asgard plot is compared with the TACK superphylum plot based on assignment of TACK genomes to arCOGs.

We employed the asCOG profiles to annotate the remaining 86 Asgard MAGs, including those that were sequenced in the later stages of this work (Supplementary Table 1). On average, 89% of the proteins encoded in these genomes were covered by asCOGs (Supplementary Table 1). Thus, the asCOGs database appears to be an efficient tool for annotation and comparative genomic analysis of Asgard MAGs and complete genomes.

### Expanding the phylogenetic diversity of Asgard archaea

Phylogenetic analysis of the Asgard MAGs based on a concatenated alignment of 209 core asCOGs (see Methods and additional data file 2) placed many of the genomes reported here into the previously delineated major Asgard lineages (Fig. 2a, Supplementary Table 1, additional data file 2), namely, Thorarchaeota (n=20), Lokiarchaeota (n=18), Hermodarchaeota (n=9), Gerdarchaeota (n=3), Helarchaeota (n=2), and Odinarchaeota (n=1). Additionally, we identified 6 previously unknown major Asgard lineages that appear to be strong candidates to become additional phyla (Fig. 2a and b, Supplementary Table 1; see also Taxonomic description of new taxa in the Supplementary Information and additional data file 1). A clade formed by As_085 and As_075 is a deeply branching sister group to the previously recognized Heimdallarchaeota(*2*). Furthermore, our phylogenetic analysis supported the further split of “Heimdallarchaeota” into 4 phylum-level lineages according to the branch length in the concatenated phylogeny (see Materials and Methods; see also Taxonomic description of new taxa in the Supplementary Information and additional data file 1). The putative phyla within the old Heimdallarchaeota included the previously defined Gerdarchaeota (*4*), and three additional phyla that could be represented by As_002 (LC2), As_003 (LC3) and AB_125 (As_001), respectively. Another 3 previously undescribed lineages were related, respectively, to Hel-, Loki-, Odin- and Thorarchaeota. Specifically, As_181, As_178 and As_183 formed a clade that was deeply rooted at the Hel-Loki-Odin-Thor clade; As_129 and As_130 formed a sister group to Odinarchaeota; and a lineage represented by As_086 was a sister group to Thorarchaeota. These results were buttressed by the 16S rRNA gene phylogeny, comparisons of the mean amino acid identity and 16S rRNA sequence identity (Fig. 2b, Supplementary Figure 3, Supplementary Figure 4, Supplementary Table 2 and Supplementary Table 3).

**Figure 2.**
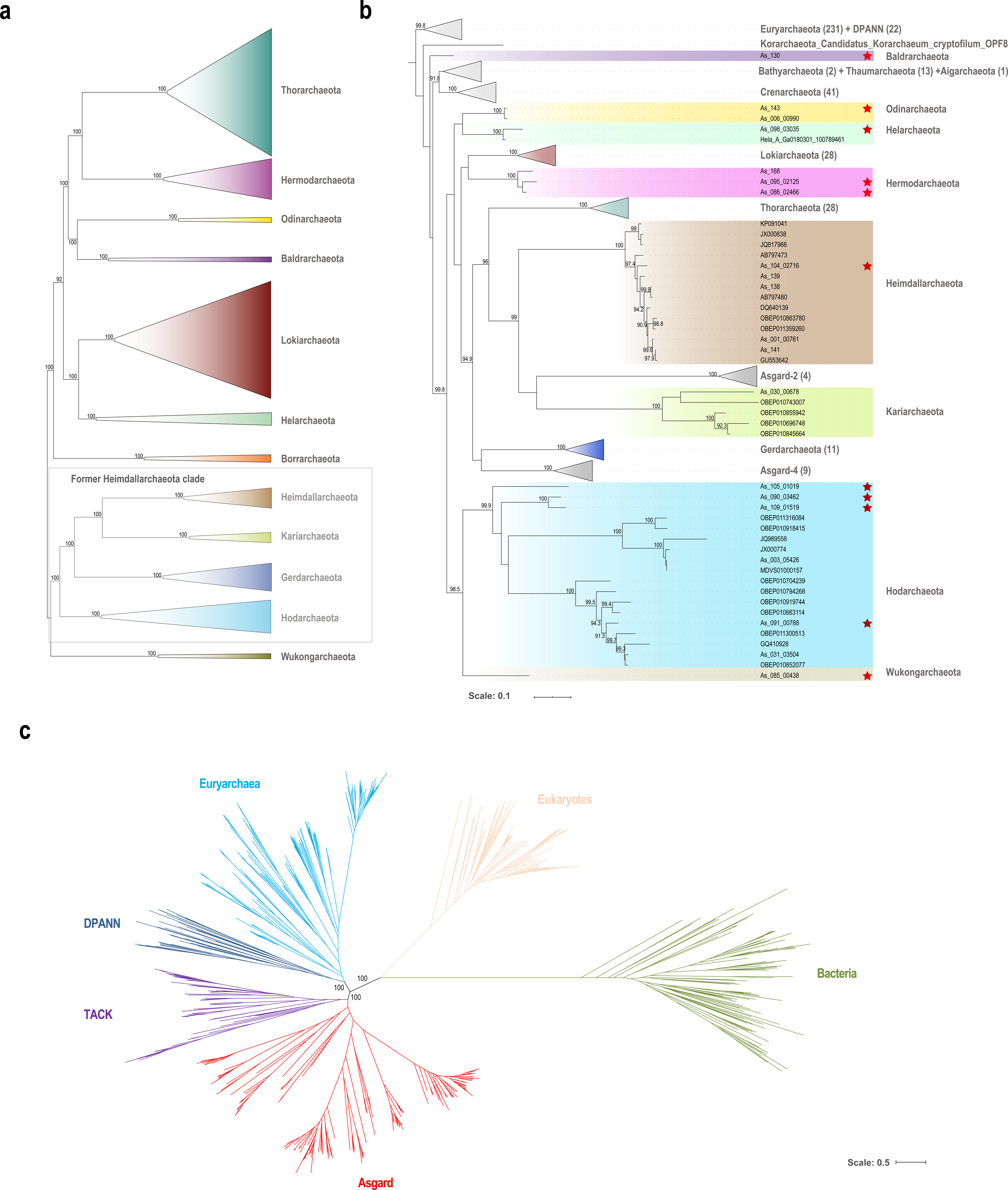
Phylogenetic analysis of Asgard archaea and their relationships with eukaryotes. **(a)** Maximum likelihood tree, inferred with IQ-tree and LG+F+R10 model, constructed from concatenated alignments of the protein sequences from 209 core Asgard Clusters of Orthologs (asCOGs). Only the 12 phylum-level clades are shown, with species within each clade collapsed. See supplementary Methods and Supplementary Table 5 for details. **(b)** Maximum likelihood tree, inferred with IQ-tree and SYM+R8 model, based on 16S rRNA gene sequences. Red stars in **(b)** denote MAGs reconstructed in the current study. **(c)** Phylogenetic tree of bacteria, archaea and eukaryotes, inferred with IQ-tree under LG+R10 model, constructed from concatenated alignments of the protein sequences of 30 universally conserved genes (see Material and Methods for details). The tree shows the relationships between the major clades. The trees are unrooted and are shown in a pseudorooted form for visualization purposes only. The actual trees and alignments are in Additional data file 2 and list of the trees are provided in the Supplementary Table 4 and 5.

We propose the name Wukongarchaeota after Wukong, a Chinese legendary figure who caused havoc in the heavenly palace, for the putative phylum represented by MAGs As_085 and As_075 (*Candidatus* Wukongarchaeum yapensis), and names of Asgard deities in the Norse mythology for the other 5 proposed phyla: (1) Hodarchaeota, after Hod, the god of darkness, for MAG As_027 (*Candidatus* Hodarchaeum mangrove); (2) Kariarchaeota, after Kari, the god of the North wind, for MAG As_030 (*Candidatus* Kariarchaeum pelagius); (3) Borrarchaeota after Borr, the creator god and father of Odin, for MAG As_133 (*Candidatus* Borrarchaeum yapensis); (4) Baldrarchaeota, after Baldr, the god of light and brother of Thor, for MAG As_130 (*Candidatus* Baldrarchaeum yapensis); (5) and Hermodarchaeota after Hermod, the messenger of the gods, son of Odin and brother of Baldr, for MAG As_086 (*Candidatus* Hermodarchaeum yapensis) (Fig. 2a). For details, see Taxonomic Description of new taxa in the **Supplementary Information**.

The gene content of Asgard MAGs agrees well with the phylogenetic structure of the group. The phyletic patterns of the asCOG form clusters that generally correspond to the clades identified by phylogenetic analysis (Fig. 3a), suggesting that gene gain and loss within Asgard archaea largely proceeded in a clock-like manner and/or that horizontal gene exchange preferentially occurred between genomes within the same clade.

**Figure 3.**
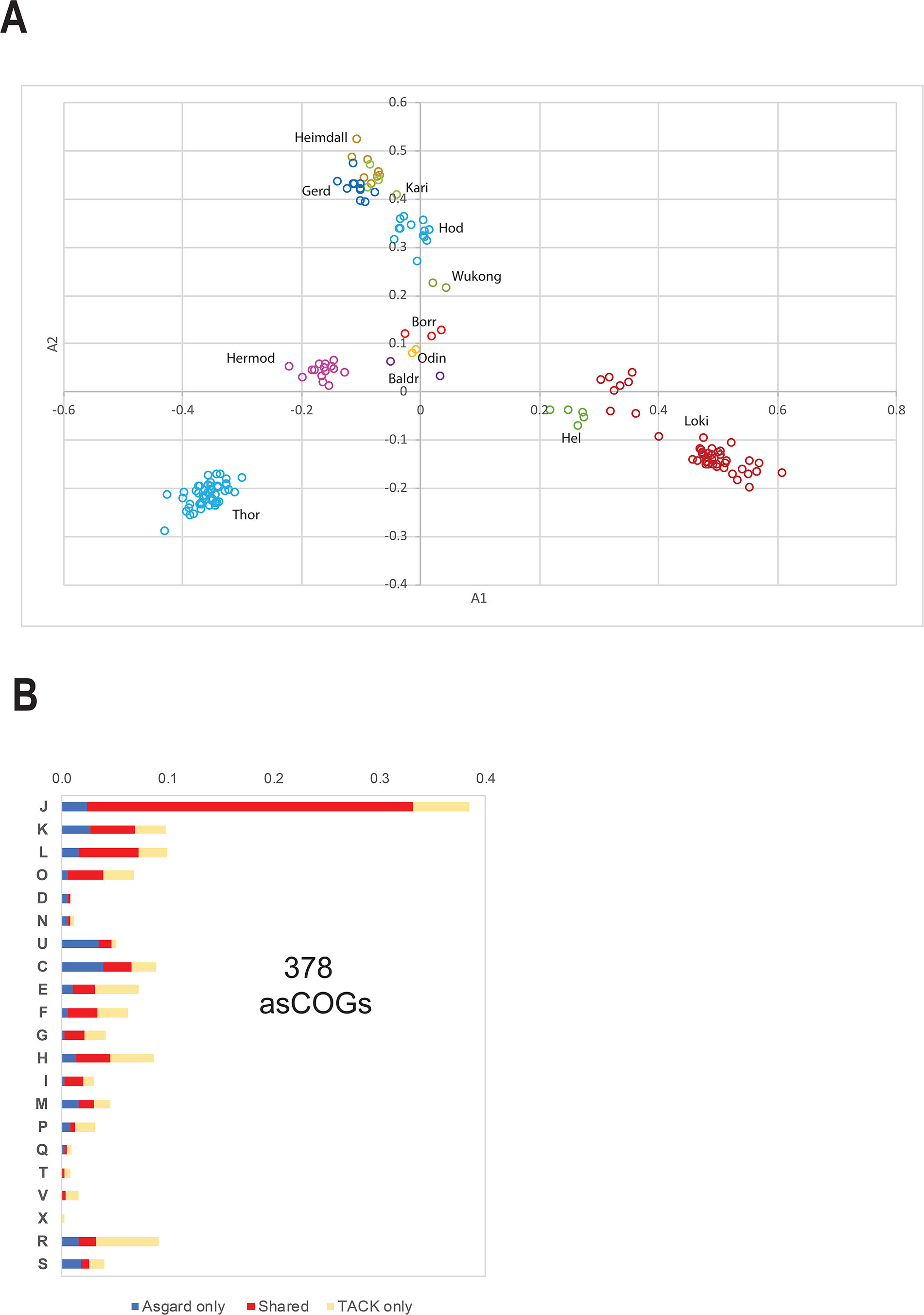
Phyletic patterns of asCOGs and functional distribution of Asgard core genes. **(a)** Classical Multidimensional Scaling analysis of binary presence-absence phyletic patterns for 13,939 asCOGs that are represented in at least two genomes (see Material and Methods for details). **(b)** Functional breakdown of Asgard core genes (378 asCOGs) compared with TACK superphylum core genes (489 arCOGs). The values were normalized as described in Materials and Methods. Functional classes of genes: J, Translation, ribosomal structure and biogenesis; K, Transcription; L, Replication, recombination and repair; D, Cell cycle control, cell division, chromosome partitioning; V, Defense mechanisms; T, Signal transduction mechanisms; M, Cell wall/membrane/envelope biogenesis; N, Cell motility; U, Intracellular trafficking, secretion, and vesicular transport; O, Posttranslational modification, protein turnover, chaperones; X, Mobilome: prophages, transposons; C, Energy production and conversion; G, Carbohydrate transport and metabolism; E, Amino acid transport and metabolism; F, Nucleotide transport and metabolism; H, Coenzyme transport and metabolism; I, Lipid transport and metabolism; P, Inorganic ion transport and metabolism; Q, Secondary metabolites biosynthesis, transport and catabolism; R, General function prediction only; S, Function unknown;

### Phylogenomic analysis and positions of Asgard archaea and eukaryotes in the tree of life

The outcome of phylogenetic reconstruction, especially, when deep branchings are involved, such as those that are relevant for the 3D vs 2D conundrum, depends on the phylogenetic methods employed, the selection of genes for phylogeny construction and, perhaps most dramatically, on the species sampling (*22*–*24*). In many cases, initially uncertain positions of lineages in a tree settle over time once more representatives of the groups in question and their relatives become available.

In our analysis of the universal phylogeny, we aimed to make the species set for phylogenetic reconstruction as broadly representative as possible, while keeping its size manageable, to allow the use of powerful phylogenetic methods. The tree was constructed from alignments of conserved proteins of 162 Asgard archaea, 286 other archaea, 98 bacteria and 72 eukaryotes (see Supplementary Material and Methods for details of the procedure including the selection of a representative species set and Supplementary Table 4). Members of 30 families of (nearly) universal proteins that appear to have evolved without much HGT and have been previously employed for the reconstruction of the tree of life (*25*) were used to generate a concatenated alignment of 7411 positions, after removing low information content positions (Supplementary Table 4). For the phylogenetic reconstruction, we used the IQ-tree program with several phylogenetic models (see Methods and Supplementary Table 5 for details). Surprisingly, the resulting trees had the 3D topology, with high support values for all key bifurcations (Fig. 2c, additional data file 2).

A full investigation of the effects of different factors, in particular, the marker gene selection, on the tree topology is beyond the scope of this work. Nonetheless, we addressed the possibility that the 3D topology resulted from the model used for the tree reconstruction and/or the species selection. To this end, we constructed 100 trees from the same alignment by randomly sampling 5 representatives of Asgard archaea, other archaea, bacteria and eukaryotes each. For these smaller sets of species, the best model identified by Williams et al. (LG+C60+G4+F) could be employed, resulting in 50 3D and 50 2D trees (Supplementary Table 5, additional data file 2). Because IQ-tree identified this model as over-specified for such a small alignment, we also tested a more restricted model (LG+C20+G4+F), obtaining 58 3D and 42 2D trees for the same set of 100 samples (Supplementary Table 5, additional data file 2).

The results of our phylogenetic analysis indicate that: 1) species sampling substantially affects the tree topology; 2) even the set of most highly conserved genes that appear to be minimally prone to HGT, yields conflicting signals for different species sets. Additional markers, less highly conserved and more prone to HGT, are unlikely to improve phylogenetic resolution and might cause systematic error. Notably, the topology of our complete phylogenetic tree (Fig. 2a) within the archaeal clade is mostly consistent with the tree obtained in a preliminary analysis of a larger set of archaeal genomes and a larger marker gene set (*26*). Taking into account these observations and the fact that we used the largest set of Asgard archaea and other archaea compared to all previous phylogenetic analyses, the appearance of the 3D topology in our tree indicates that the origin of eukaryotes from within Asgard cannot be considered a settled issue. Various factors affecting the tree topology, including further increased species representation, particularly, of Asgard and the deep branches of the TACK superphylum, such as Bathyarchaeota and Korarchaeota, remain to be explored in order to definitively resolve the evolutionary relationship between archaea and eukaryotes.

### The core gene set of Asgard archaea

We next analyzed the core set of conserved Asgard genes which we arbitrarily defined as all asCOGs that are present at least in one third of the MAGs in each of the 12 phylum-level lineages, with the mean representation across lineages >75%. Under these criteria, the Asgard core includes 378 asCOGs (Supplementary Table 6). As expected, most of these protein families, 293 (77%), are universal (present in bacteria, other archaea and eukaryotes), 62 (16%) are represented in other archaea and eukaryotes, but not in bacteria, 15 (4%) are found in other archaea and bacteria, but not in eukaryotes, 7 (2%) are archaea-specific, and only 1 (0.003%) is shared exclusively with eukaryotes (Supplementary Figure 5). Most of the core asCOGs show comparable levels of similarity to homologs from two or all three domains of life. The second largest fraction of the core asCOGs shows substantially greater sequence similarity (at least, 25% higher similarity score) to homologous proteins from archaea than to those from eukaryotes and/or bacteria (Supplementary Table 6). Compared with the 219 genes that comprise the pan-archaeal core (*27*), the Asgard core set lacks 12 genes, each of which, however, is present in some subset of the Asgard genomes. These include three genes of diphthamide biosynthesis and 2 ribosomal proteins, L40E and L37E. The intricate evolutionary history of gene encoding translation elongation factors and enzymes of diphthamide biosynthesis in Asgard has been analyzed previously (*28*). Also of note is the displacement of the typical archaeal glyceraldehyde-3-phosphate dehydrogenase (type II) by a bacterial one (type I) in most of the Asgard genomes (cog.001204, additional data file 1).

Functional distribution of the core asCOGs is shown in Fig. 3b (also see additional data file 1). For comparison, we also derived an extended gene core for the TACK superphylum, using similar criteria (at least 50% in each of the 6 lineages and 75% of the genomes overall, Fig 3b). For at least half of the Asgard core genes, across most functional classes, there were no orthologs in the TACK core. The most pronounced differences were found, as expected, in the category U (intracellular trafficking, secretion, and vesicular transport). In Asgard archaea, this category includes 19 core genes compared with 7 genes in TACK; 13 of these genes are specific to the Asgard archaea and include components of ESCRT I and II, 3 distinct Roadblock/longin families, 2 distinct families of small GTPases, and a few other genes implicated in related processes (Supplementary Table 6).

We compared the protein annotation obtained using asCOGs with the available annotation of ‘*Candidatus* Prometheoarchaeum syntrophicum’ and found that using asCOGs allowed at least a general functional prediction for 649 of the 1756 (37%) ‘hypothetical proteins’ in this organism, the only one in Asgard with a closed genome. We also identified 139 proteins, in addition to the 80 described originally, that can be considered Eukaryotic Signature Proteins, or ESPs (see next section).

### Eukaryotic features of Asgard archaea gleaned from genome analysis

The enrichment of Asgard proteomes with homologs of eukaryote signature proteins (ESPs), such as ESCRTs, components of protein sorting complexes including coat proteins, complete ubiquitin machinery, actins and actin-binding proteins gelsolins and profilins, might be the strongest argument in support of a direct evolutionary relationship between Asgard archaea and eukaryotes (*2*, *29*). However, the definition of ESPs is fuzzy because many of these proteins, in addition to their occurrence in Asgard, are either scattered among several other archaeal genomes, often, from diverse groups (*30*), or consist of promiscuous domains that are common in archaea, bacteria and eukaryotes, such as WD40 (after the conserved terminal amino acids of the repeat units, also known as beta-transducin repeats), LRR (Leucine-Rich Repeats), TPR (TetratricoPeptide Repeats), HEAT (Huntingtin-EF3-protein phosphatase 2A-TOR1) and other, largely, repetitive domains (*31*, *32*). Furthermore, the sequences of some ESPs have diverged to the extent that they become hardly detectable with standard computational methods. Our computational strategy for delineating an extensive yet robust ESP set is described under Materials and Methods. The ESP set we identified contained 505 asCOGs, including 238 that were not closely similar (E-value=10^−10^, length coverage 75%) to those previously described by Zaremba-Niedzwiedzka et al. (*2*)(Supplementary Table 7). In a general agreement with previous observations, the majority of these ESPs, 329 of the 505, belonged to the ‘Intracellular trafficking, secretion, and vesicular transport’ (U) functional class, followed by ‘Posttranslational modification, protein turnover, chaperones’ (O), with 101 asCOGs (Supplementary Table 7). Among the asCOGs in the U class, 130 were Roadblock/LC7 superfamily proteins, including longins, sybindin and profilins, and 94 were small GTPases of several families, such as RagA-like, Arf-like and Rab-like ones, as discussed previously (*33*).

The phyletic patterns of ESP asCOGs in Asgard archaea are extremely patchy and largely lineage-specific (Fig. 4), indicating that most of the proteins in this set are not uniformly conserved throughout Asgard evolution, but rather, are prone to frequent HGT, gene losses and duplications. These evolutionary processes are correlated in prokaryotes, resulting in the overall picture of highly dynamic evolution (*34*). Even the most highly conserved ESP asCOG are missing in some Asgard lineages but show multiple duplications in others (Fig. 4 and Supplementary Table 7). Surprisingly, many gaps in the ESPs distribution were detected in the Heimdallarchaeota that include the suspected ancestors of eukaryotes.

**Figure 4.**
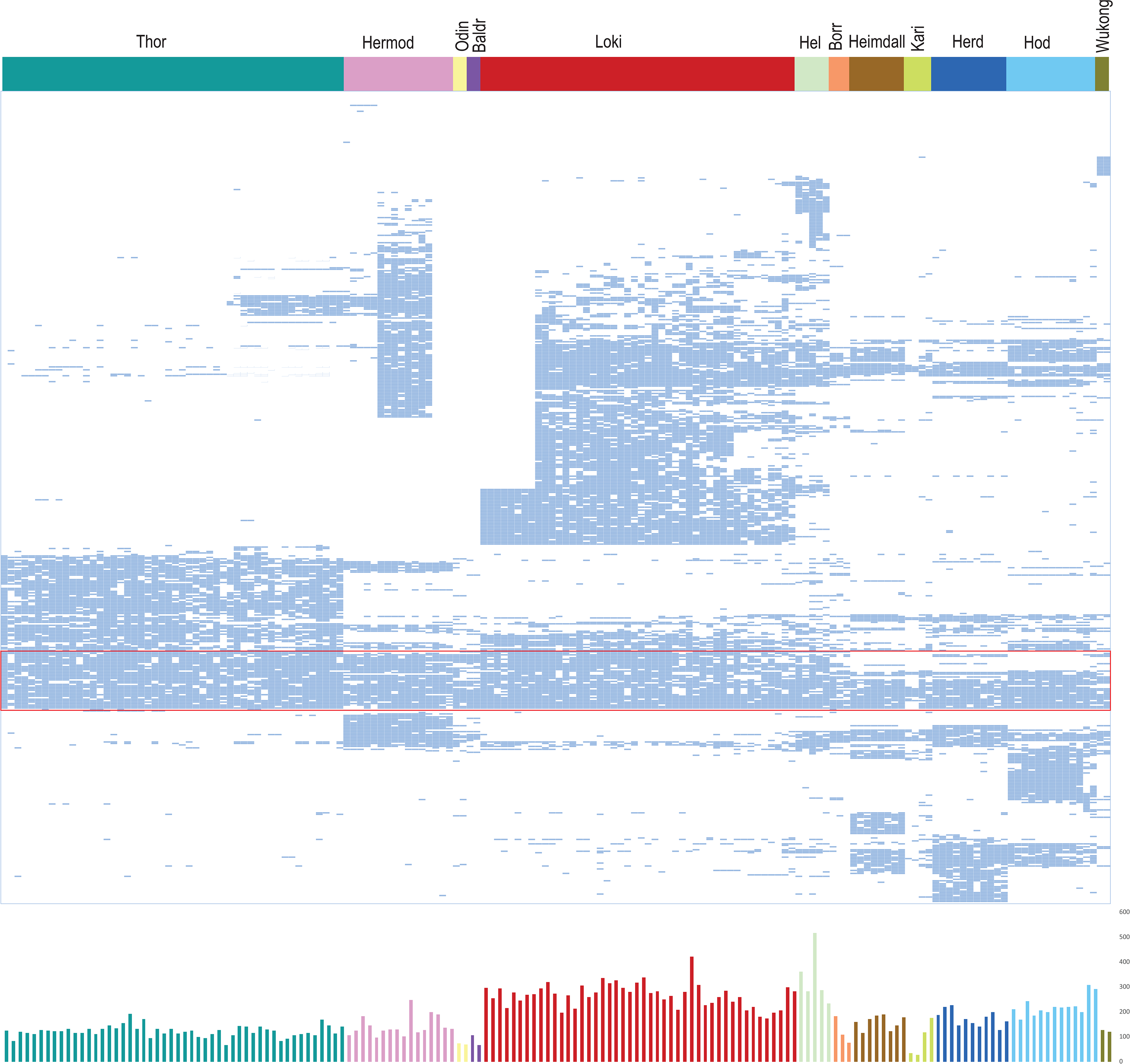
Phyletic patterns of Eukaryotic Signature Proteins (ESPs) encoded in Asgard genomes. All 505 ESP asCOGs are grouped by distance between binary presence-absence phyletic patterns. The most highly conserved ESP asCOGs are shown within the red rectangle. Below the plot of the number of ESP domains in each genome is shown. For details, see Supplementary Table 7.

Characteristically, many ESPs are multidomain proteins, with 37% assigned to more than one asCOG, compared to 17% among non-ESP proteins (Supplementary Table 7). Some multidomain ESPs in Asgard archaea have the same domain organizations as their homologs in eukaryotes, but these are a minority and typically contain only two domains. Examples include the fusion of two EAP30/Vps37 domains (*35*), and Vps23 and E2 domains (*35*) in ESCRT complexes, multiple Rag family GTPases, in which longin domain is fused to the GTPase domain, and several others. By contrast, most of the domain architectures of the multidomain ESP proteins were not detected in eukaryotes and often are found only in a narrow subset of Asgard archaea, suggesting extensive domain shuffling during Asgard evolution (Fig. 5a). For example, we identified many proteins containing a fusion of Vps28/Vps23 from ESCRT I complex (*35*) with C-terminal domains of several homologous subunits of adaptin and COPI coatomer complexes (*36*, *37*), and E3 UFM1-protein ligase 1, which is involved in the UFM1 ubiquitin pathway (*38*) (Fig. 5a). Generally, a protein with such a combination of domains can be predicted to be involved in ubiquitin-dependent membrane remodeling but, because its domain architecture is unique, the precise function cannot be inferred.

**Figure 5.**
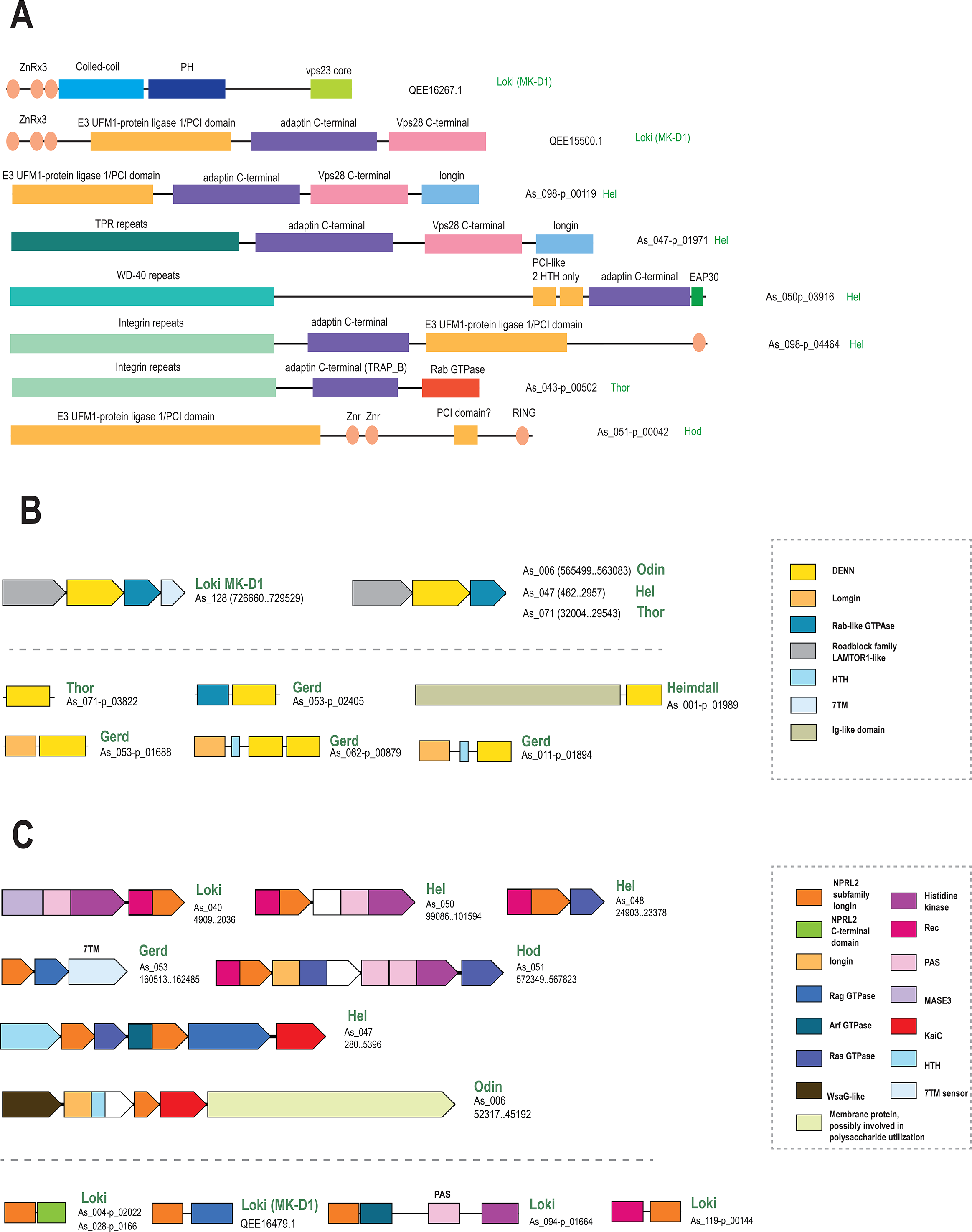
Domain architectures of selected Asgard ESPs. **(a) ESPs with unique domain architectures.** The schematic of each multidomain protein is roughly proportional to the respective protein length. The identified domains are shown inside the arrows approximately according to their location and are briefly annotated. Homologous domains are shown by the same color or pattern. **(b) DENN domain proteins in Asgards.** Upper part of the figure above the dashed line shows putative operon encoding DENN domain proteins. Genes are shown by block arrows with the length proportional to the size of the corresponding protein. For each protein, the nucleotide contig or genome partition accession number, Asgard genome ID and lineage are indicated. The part of the figure below the dashed line shows domain organization of diverse proteins containing DENN domain. Homologous domains are shown by the same color. The inset on the right side explains identity of the domains **(c) NPRL2-like proteins in Asgards.** Designations are the same as in Figure 4a.

The majority of the ESP genes of Asgard archaea do not belong to conserved genomic neighborhoods, but several such putative operons were detected. Perhaps, the most notable one is the ESCRT neighborhood which includes genes coding for subunits of ESCRT I, II and III, and often, components of the ubiquitin system (*2*), suggesting an ancient link between the two systems that persists in eukaryotes (*35*). We predicted another operon that is conserved in most Asgard archaea and consists of genes encoding a LAMTOR1-like protein of the Roadblock superfamily, a Rab-like small GTPase, and a protein containing the DENN (differentially expressed in normal and neoplastic cells) domain that so far has been identified only in eukaryotes (Fig. 5b). Two proteins consisting of a DENN domain fused to longin are subunits of the folliculin (FLCN) complex that is conserved in eukaryotes. The FLCN complex is the sensor of amino acid starvation interacting with Rag GTPase and Ragulator lysosomal complex, and a key component of the mTORC1 pathway, the central regulator of cell growth in eukaryotes (*39*). Some Heimdallarchaea encode several proteins with the exact same domain organization as FLCN (Fig. 5b). Ragulator is a complex that consists of 5 subunits, each containing the Roadblock domain. In Asgard archaea, however, the GTPase present in the operon is from a family that is distinct from the Rag GTPases, which interact with both FLCN and Ragulator complexes in eukaryotes, despite the fact that Rag family GTPases are abundant in Asgards (*33*) (Supplementary Table 7). Nevertheless, this conserved module of Asgard proteins is a strong candidate to function as a guanine nucleotide exchange factor for Rab and Rag GTPases, analogously to the eukaryotic FLCN. In eukaryotes, the DENN domain is present in many proteins with different domain architectures that interact with different partners and perform a variety of functions (*40*, *41*). The Asgard archaea also encode other DENN domain proteins, and the respective genes form expanded families of paralogs in Loki, Hel and Heimdall lineages, again, with domain architectures distinct from those in eukaryotes (Fig. 5b) (*42*).

Prompted by the identification of a FLCN-like complex, we searched for other components of the mTORC1 regulatory pathway in Asgard archaea. The GATOR1 complex that consists of three subunits, Depdc5, Nprl2, and Nprl3, is another amino acid starvation sensor that is involved in this pathway in eukaryotes (*43*). Nitrogen permease regulators 2 and 3 (NPRL2 and NPRL3) are homologous GATOR1 subunits that contain a longin domain and a small NPRL2-specific C-terminal domain (*43*). We identified a protein family with this domain organization in most Thor MAGs and a few Loki MAGs. Several other ESP asCOGs include proteins with high similarity to the longin domain of NPRL2. Additionally, we identified many fusions of the NPRL2-like longin domain with various domains related to prokaryotic two-component signal transduction system (Fig. 5c). Considering the absence of a homolog of phosphatidylinositol 3-kinase, the catalytic domain of the mTOR protein, it seems likely that, in Asgard archaea, the key growth regulation pathway remains centered at typical prokaryotic two-component signal transduction systems whereas at least some of the regulators and sensors in this pathway are “eukaryotic”. The abundance of NPRL2-like longin domains in Asgard archaea implies that the link between this domain and amino acid starvation regulation emerged at the onset of Asgard evolution if not earlier.

### Diverse metabolic repertoires, ancestral metabolism of Asgard archaea, and syntrophic evolution

Examination of the distribution of the asCOGs among the 12 Asgard archaeal phyla showed that the metabolic pathway repertoire was conserved among the MAGs of each phylum but differed between the phyla (Fig. 3a). Three distinct lifestyles were predicted by the asCOG analysis for different major branches of Asgard archaea, namely, anaerobic heterotrophy, facultative aerobic heterotrophy, and chemolithotrophy (Fig. 6, Supplementary Figure 9). For the last Asgard archaeal common ancestor (LAsCA), a mixotrophic life style, including both production and consumption of H_2_, can be inferred from parsimony considerations (Fig. 6, Supplementary Table 8; see Materials and Methods for further details). Loki-, Thor-, Hermod-, Baldr- and Borrarchaeota encode all enzymes for the complete (archaeal) Wood-Ljungdahl pathway (WLP) and are predicted to oxidize organic substrates, likely, by using the reverse WLP, given the lack of enzymes for oxidation of inorganic compounds (e.g., hydrogen, sulfur/sulfide and nitrogen/ammonia). The genomes of these five Asgard phyla encode homologues of membrane-bound respiratory H_2_-evolving Group 4 [NiFe] hydrogenase (Supplementary Figure 6) and/or cytosolic cofactor-coupled bidirectional Group 3 [NiFe] hydrogenase (*44*) (Supplementary Figure 7). Phylogenetic analysis of both group 3 and group 4 [NiFe] hydrogenases showed that Asgard archaea form distinct clades well separated from the functionally characterized hydrogenases, hampering the prediction of their specific functions in Asgard archaea. The functionally characterized group 4 [NiFe] hydrogenases in the Thermococci are involved in the fermentation of organic substrates to H_2_, acetate and carbon dioxide (*45*, *46*). The presence of group 3 [NiFe] hydrogenases suggests that these Asgard archaea cannot use H_2_ as an electron donor because they lack the enzyme complex coupling H_2_ oxidation to membrane potential generation. Thus, in these organisms, bifurcate electrons from H_2_ are likely to be used to support the fermentation of organic substrates exclusively (*45*–*47*).

**Figure 6.**
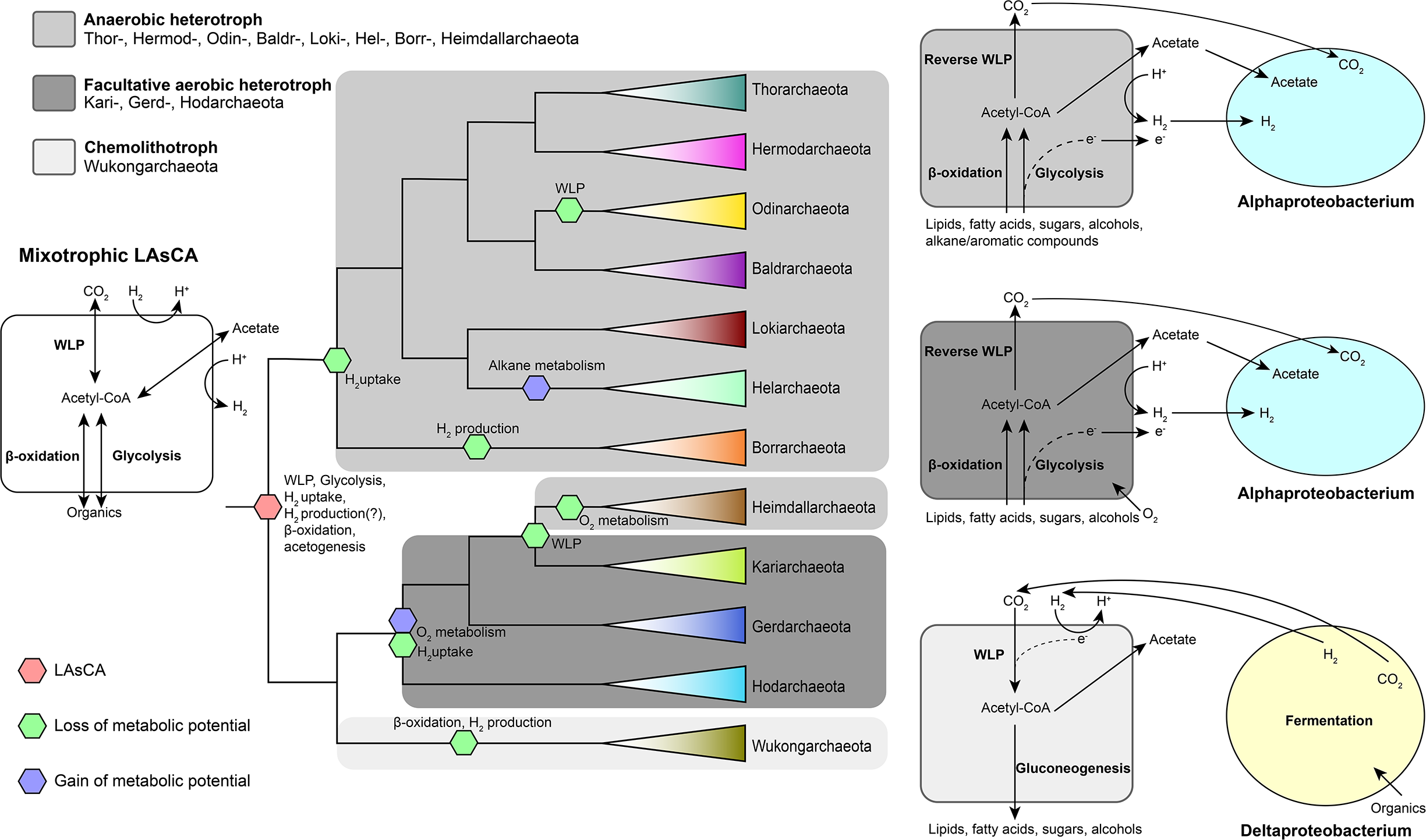
Reconstruction and evolution of the key metabolic processes in Asgard archaea. The schematic phylogeny of Asgard archaea is from Figure 1a. LAsCA, Last Asgard Common Ancestor; WLP, Wood-Ljungdahl pathway.

Both Wukongarchaeota genomes (As_075 and As_085) encode a bona fide membrane-bound Group 1k [NiFe] hydrogenase that could mediate hydrogenotrophic respiration using heterodisulfide as the terminal electron acceptor (*48*, *49*) (Fig. 6, Supplementary Figure 8, Supplementary Figure 10). The group 1k [NiFe] hydrogenase is exclusively found in methanogens of the order Methanosarcinales (Euryarchaoeta) (*50*), and it is the first discovery of the group 1 [NiFe] hydrogenase in the Asgard archaea. Wukongarchaeota also encode all enzymes for a complete WLP and a putative ADP-dependent acetyl-CoA synthetase for acetate synthesis. Unlike all other Asgard archaea, Wukongarchaeota lack genes for citrate cycle and beta-oxidation. Thus, Wukongarchaeota appear to be obligate chemolithotrophic acetogens. The genomes of Wukongarchaeota were discovered only in seawater of the euphotic zone of the Yap trench (0 m and 125 m). Dissolved H_2_ concentration is known to be the highest in surface seawater, where the active microbial fermentation, compared to deep sea (*51*), could produce sufficient amounts of hydrogen for the growth of Wukongarchaeota. Hodarchaeota, Gerdarchaeota, Kariarchaeota, and Heimdallarchaeota share a common ancestor with Wukongarchaeota (Fig. 6). However, genome analysis implies different lifestyles for these organisms. Hod-, Gerd- and Kariarchaeota encode various electron transport chain components, including heme/copper-type cytochrome/quinol oxidase, nitrate reductase, and NADH dehydrogenase, most likely, allowing the use of oxygen and nitrate as electron acceptors during aerobic and anaerobic respiration, respectively (*44*). In addition, Hod-, Gerd- and Heimdallarchaeota encode phosphoadenosine phosphosulfate (PAPS) reductase and adenylylsulfate kinase for sulfate reduction, enabling the use of sulfate as electron acceptor during anaerobic respiration. Gerd-, Heimdall-, and Hodarchaeota are only found in coastal and deep-sea sedimentary environments, whereas Kariarchaeota were found also in marine water. The versatile predicted metabolic capacities of these groups suggest that Hod-, Gerd- and Kariarchaeota might occupy both anoxic and oxic niches. In contrast, Heimdallarchaeota appear to be able to thrive only in anoxic environments.

In the widely considered syntrophy scenarios (*52*), eukaryogenesis has been proposed to involve metabolic symbiosis (syntrophy) between an archaeon and one or two bacterial partners which, in the original hydrogen/syntrophy hypothesis, were postulated to donate H_2_ for methane or hydrogen sulfide production by the consortium (*53*, *54*). The syntrophic scenarios were boosted by the discovery of apparent syntrophy between *Candidatus* P. syntrophicum and Deltaproteobacteria which led to the proposal of the Etangle-Engulf-Endogenize (E^3^) model of eukaryogenesis (*17*, *55*). Reconstruction of the Lokiarchaeon metabolism has suggested that this organism was hydrogen-dependent, in accord with the hydrogen-syntrophic scenarios (*54*). In contrast, subsequent analysis of the metabolic potentials of 4 Asgard phyla has led to the inference that these organisms were primarily organoheterotrophic and H_2-_ producing, the ‘reverse flow model’ model of protoeukaryote energy metabolism that involves electron or hydrogen flow from an Asgard archaeon to the alphaproteobacterial ancestor of mitochondria, in the opposite direction from that in the original hydrogen-syntrophy hypotheses (*44*). Here, we discovered a deeply branching Asgard group, Wukongarchaeota, which appears to include obligate hydrogenotrophic acetogens, suggesting the possibility of the LAsCA being a hydrogen-dependent autotroph (Fig. 6). This finding suggests that LAsCA both produced and consumed H_2_. Thus, depending on the exact relationship between Asgard archaea and eukaryotes that remains to be elucidated, our findings could be compatible with different syntrophic scenarios that postulate H_2_ transfer from bacteria to the archaeal symbiont or in the opposite direction.

## Conclusions

The Asgard archaea that were discovered only 5 years ago as a result of the painstaking assembly of several Loki Castle metagenomes have grown into a highly diverse archaeal superphylum. The most remarkable feature of the Asgards is their apparent evolutionary affinity with eukaryotes that has been buttressed by two independent lines of evidence: phylogenetic analysis of highly conserved genes and detection of multiple ESPs that are absent or far less common in other archaea. The 75 MAGs added here substantially expand the phylogenetic and metabolic diversity of the Asgard superphylum. The extended set of Asgard genomes provides for a phylogeny based on a far more representative species sampling than available previously and a substantially expanded ESP analysis employing powerful computational methods. This extended analysis reveals 6 putative additional Asgard phyla but does not immediately clarify the Asgard-eukaryote relationship. Indeed, phylogenetic analysis of conserved genes in an expanded set of archaea, bacteria and eukaryotes yields conflicting 3D and 2D signals, with an unexpected preference for the 3D topology. Thus, the conclusion that eukaryotes emerged from within Asgard archaea, in particular, from the Heimdall lineage, appears to be premature. Further phylogenomic study with an even broader representation of diverse archaeal lineages as well as, possibly, even more sophisticated evolutionary models are required to clarify the relationships between archaea and eukaryotes.

Our analysis of Asgard genomes substantially expanded the set of ESPs encoded by this group of archaea and revealed numerous, complex domain architectures of these proteins. These results further emphasize the excess of ESPs in Asgards compared to other archaea and provide additional support to the conclusion that most of the Asgard ESPs are involved in membrane remodeling and intracellular trafficking. However, in parallel with the phylogenomic results, detailed analysis of the ESPs reveals a complex picture. Most of the multidomain Asgard ESPs possess domain architectures distinct from typical eukaryotic ones and some of these arrangements include signature prokaryotic domains, suggesting substantial functional differences from the respective eukaryotic systems. Furthermore, virtually all the ESPs show patchy distributions in Asgard and other archaea, indicative of a history of extensive HGT, gene losses and paralogous family expansion. All these findings seem to be best compatible with the model of a dispersed, dynamic archaeal ‘eukaryome’ (*30*) that widely spreads among archaea via HGT, so far reaching the highest ESP density in the Asgard archaea.

The results of this work cannot rule out the possibility of the emergence of eukaryotes from within the Asgard but seem to be better compatible with a different evolutionary scenario under which the conserved core of eukaryote genes involved in informational processes originates from an as yet unknown ancestor group that might be a deep archaeal branch or could lie outside the presently characterized archaeal diversity. These hypothetical ancestral forms might have accumulated components of the mobile archaeal ‘eukaryome’ to an even greater extent than the Asgards archaea, eventually, giving rise to eukaryote-like cells, likely, via a form of syntrophy with one or more bacterial partners. Combined genomic and (undoubtedly, far more challenging) biological study of diverse archaea is essential for further advancing our understanding of eukaryogenesis.

## Author contributions

ML, EVK, KSM and YL initiated the study;; YL performed metagenomic assembly, binning, metabolism analysis; KSM, AN and YIW performed comparative genomic analysis; YL, KSM, YIW and WCH performed phylogenetic analysis; KSM and YIW constructed asCOGs; KSM, YIW, YL, ML, and EVK analyzed the data; YL, KSM, WCH, EVK, and ML wrote the manuscript that was read, edited and approved by all authors.

## Acknowledgements

ML and YL are supported by National Natural Science Foundation of China (Grant No. 91851105, 31970105 and 31700430), the Key Project of Department of Education of Guangdong Province (No.2017KZDXM071), and the Shenzhen Science and Technology Program (Grant no. JCYJ20170818091727570 and KQTD20180412181334790). KSM, YIW, SAS and EVK are supported by the Intramural Research Program of the National Institutes of Health of the USA (National Library of Medicine).

## Supplementary Information

### Taxonomic Description of new taxa

#### *Candidatus* Wukongarchaeum

(Wu.kong.ar.chae’um. N. L. n. Wukong a legendary Chinese figure, also known as Monkey King, who caused havoc in the heavenly palace); N.L. neut. N. *archaeum* (from Gr. adj. archaios ancient) archaeon; N. L. neut. N. Wukongarchaeum.

#### *Candidatus* Wukongarchaeum yapensis

(yap’ensis N. L. masc. adj. pertaining to Yap trench, which is the geographical position where the first type material of this species was obtained). Type material is the genome designated as As_085 (Yap4.bin4.70) representing ‘*Candidatus* Wukongarchaeum yapensis’. The genome “As_085” represents a MAG consisting of 2.16 Mbps in 277 contigs with an estimated completeness of 92.52%, an estimated contamination of 4.05%, a 16S and 23S rRNA gene and 14 tRNAs. The MAG recovered from a marine water metagenome (Yap trench, Western Pacific), with an estimated depth of coverage of 31.4, has a GC content of 38%.

#### *Candidatus* Hodarchaeum

(Hod.ar.chae’um. N. L. n. Hod a son of Odin in Norse mythology); N.L. neut. N. archaeum (from Gr. adj. archaios ancient) archaeon; N. L. neut. N. Hodarchaeum.

#### *Candidatus* Hodarchaeum mangrovi

(man.gro’vi N.L. fem. n. of a mangrove, referring to the isolation of the type material from mangrove soil). Type material is the genome designated as As_027 (FT2_5_011) representing ‘*Candidatus* Hodarchaeum mangrovi’. The genome “As_027” represents a MAG consisting of 4.01 Mbps in 348 contigs with an estimated completeness of 93.61%, an estimated contamination of 0.93%, a 23S rRNA gene and 14 tRNAs. The MAG recovered from mangrove sediment metagenomes (Futian Nature Reserve, China), with an estimated depth of coverage of 17.9, has a GC content of 32.9%.

#### *Candidatus* Kariarchaeum

(Ka.ri.ar.chae’um. N. L. n. Kari the god of wind and brother to Aegir in Norse mythology); N.L. neut. N. *archaeum* (from Gr. adj. archaios ancient) archaeon; N. L. neut. N. Kariarchaeum.

#### *Candidatus* Kariarchaeum pelagius

(pe.la’gi.us. L. masc. adj. of or belonging to the sea, referring to the isolation of the type material from the Ocean). Type material is the genome designated as As_030 (RS678) representing ‘*Candidatus* Kariarchaeum pelagius’. The genome “As_030” represents a MAG consisting of 1.41 Mbps in 76 contigs, an estimated completeness of 83.18%, with an estimated contamination of 1.87%, a 23S, 16S and 5S rRNA genes and 18 tRNAs. The MAG recovered from a marine metagenome (Saudi Arabia: Red Sea) has a GC content of 30.11%.

#### *Candidatus* Borrarchaeum

(Borr.ar.chae’um. N. L. n. Borr a creator god and father of Odin); N.L. neut. N. *archaeum* (from Gr. adj. archaios ancient) archaeon; N. L. neut. N. Borrarchaeum.

#### *Candidatus* Borrarchaeum yapensis

(yap’ensis N. L. masc. adj. pertaining to Yap trench, which is the geographical position where the first type material of this species was obtained). Type material is the genome designated as As_181 (Yap2000.bin9.141) representing ‘*Candidatus* Borrarchaeum yapensis’. The genome “As_181” represents a MAG consisting of 3.63 Mbps in 125 contigs, with an estimated completeness of 95.02%, an estimated contamination of 5.61% and 11 tRNAs. The MAG, recovered from a marine water metagenome (Yap trench, Western Pacific) with an estimated depth coverage of 15.04, has a GC content of 37.1%.

#### *Candidatus* Baldrarchaeum

(Bal.dr.ar.chae’um. N. L. n. Baldr the god of light and son of Odin and borther of Thor in Norse mythology); N.L. neut. N. *archaeum* (from Gr. adj. archaios ancient) archaeon; N. L. neut. N. Baldrarchaeum.

#### *Candidatus* Baldrarchaeum yapensis

(yap’ensis N. L. masc. adj. pertaining to Yap trench, which is the geographical position where the first type material of this species was obtained). Type material is the genome designated as As_130 (Yap30.bin9.72) representing ‘*Candidatus* Baldrarchaeum yapensis’. The genome “As_130” represents a MAG consisting of 2.27 Mbps in 100 contigs, with an estimated completeness of 93.93%, an estimated contamination of 3.74%, a 23S and 16S rRNA gene and 15 tRNAs. The MAG, recovered from a marine water metagenome (Yap trench, Western Pacific) with an estimated depth coverage of 39.99, has a GC content of 45.95%.

#### *Candidatus* Hermodarchaeum

(Her.mod.ar.chae’um. N. L. n. Hermod, messengers of the gods in the Norse mythology and son of Odin and brother of Baldr in the Norse mythology); N.L. neut. N. *archaeum* (from Gr. adj. archaios ancient) archaeon; N. L. neut. N. Hermodarchaeum.

#### *Candidatus* Hermodarchaeum yapensis

(yap’ensis N. L. masc. adj. pertaining to Yap trench, which is the geographical position where the first type material of this species was obtained). Type material is the genome designated as As_086 (Yap4.bin9.105) representing ‘*Candidatus* Hermodarchaeum yapensis’. The genome ‘As_086’ represent a MAG consisting of 2.71 Mbps in 77 contigs, with an estimated completeness of 92.99%, an estimated contamination of 1.87%, a 23S and 16S rRNA gene and 16 tRNAs. The MAG, recovered from a marine water metagenome (Yap trench, Western Pacific) with an estimated depth coverage of 19.24, has a GC content of 44.69%.

#### *Candidatus* Wukongarchaeaceae

(Wu.kong.ar.chae.a.ce’ae. N.L. neut. n. Wukongarchaeum a (Candidatus) type genus of the family; -aceae ending to denote the family; N.L. fem. pl. n. Wukongarchaeaceae the Wukongarchaeum family).

The family is delineated based on 209 concatenated Asgard Cluster of Orthologs (AsCOGs) and 16S rRNA gene phylogeny. The description is the same as that of its sole genus and species. Type genus is *Candidatus* Wukongarchaeum.

#### *Candidatus* Wukongarchaeales

(Wu.kong.ar.chae.a’les. N.L. neut. n. Wukongarchaeum a (Candidatus) type genus of the order; -ales ending to denote the order; N.L fem. pl. n. Wukongarchaeales the Wukongarchaeum order).

The order is delineated based on 209 concatenated AsCOGs and 16S rRNA gene phylogeny. The description is the same as that of its sole genus and species. Type genus is Candidatus Wukongarchaeum.

#### *Candidatus* Wukongarchaeia

(Wu.kong.ar.chae’i.a. N.L. neut. n. Wukongarchaeum a (Candidatus) type genus of the order of the class; -ia ending to denote the class; N.L fem. pl. n. Wukongarchaeia the Wukongarchaeum class).

The class is delineated based on 209 concatenated AsCOGs and 16S rRNA gene phylogeny. The description is the same as that of its sole and type order *Candidatus* Wukongarchaeales.

#### *Candidatus* Wukongarchaeota

(Wu.kong.ar.chae.o’ta. N.L. neut. n. Wukongarchaeum a (Candidatus) type genus of the class of the phylum; -ota ending to denote the phylum; N.L neut. pl. n. Wukongarchaeota the Wukongarchaeum phylum)

The phylum is delineated based on 209 concatenated AsCOGs and 16S rRNA gene phylogeny. The description is the same as that of its sole and type class *Candidatus* Wukongarchaeia.

#### *Candidatus* Hodarchaeaceae

(Hod.ar.chae.a.ce’ae. N.L. neut. n. Hodarchaeum a (Candidatus) type genus of the family; -aceae ending to denote the family; N.L. fem. pl. n. Hodarchaeaceae the Hodarchaeum family).

The family is delineated based on 209 concatenated AsCOGs and 16S rRNA gene phylogeny. The description is the same as that of its sole genus and species. Type genus is *Candidatus* Hodarchaeum.

#### *Candidatus* Hodarchaeales

(Hod.ar.chae.a’les. N.L. neut. n. Hodarchaeum a (Candidatus) type genus of the order; -ales ending to denote the order; N.L fem. pl. n. Hodarchaeales the Hodarchaeum order).

The order is delineated based on 209 concatenated AsCOGs and 16S rRNA gene phylogeny. The description is the same as that of its sole genus and species. Type genus is *Candidatus* Hodarchaeum.

#### *Candidatus* Hodarchaeia

(Hod.ar.chae’i.a. N.L. neut. n. Hodarchaeum a (Candidatus) type genus of the order of the class; -ia ending to denote the class; N.L fem. pl. n. Hodarchaeia the Hodarchaeum class).

The class is delineated based on 209 concatenated AsCOGs and 16S rRNA gene phylogeny. The description is the same as that of its sole and type order *Candidatus* Hodarchaeales.

#### *Candidatus* Hodarchaeota

(Hod.ar.chae.o’ta. N.L. neut. n. Hodarchaeum a (Candidatus) type genus of the class of the phylum; -ota ending to denote the phylum; N.L neut. pl. n. Hodarchaeota the Hodarchaeum phylum)

The phylum is delineated based on 209 concatenated AsCOGs and 16S rRNA gene phylogeny. The description is the same as that of its sole and type class *Candidatus* Hodarchaeia.

#### *Candidatus* Hodarchaeaceae

(Hod.ar.chae.a.ce’ae. N.L. neut. n. Hodarchaeum a (Candidatus) type genus of the family; -aceae ending to denote the family; N.L. fem. pl. n. Hodarchaeaceae the Hodarchaeum family).

The family is delineated based on 209 concatenated AsCOGs and 16S rRNA gene phylogeny. The description is the same as that of its sole genus and species. Type genus is *Candidatus* Hodarchaeum.

#### *Candidatus* Hodarchaeales

(Hod.ar.chae.a’les. N.L. neut. n. Hodarchaeum a (Candidatus) type genus of the order; -ales ending to denote the order; N.L fem. pl. n. Hodarchaeales the Hodarchaeum order).

The order is delineated based on 209 concatenated AsCOGs and 16S rRNA gene phylogeny. The description is the same as that of its sole genus and species. Type genus is *Candidatus* Hodarchaeum.

#### *Candidatus* Hodarchaeia

(Hod.ar.chae’i.a. N.L. neut. n. Hodarchaeum a (Candidatus) type genus of the order of the class; -ia ending to denote the class; N.L fem. pl. n. Hodarchaeia the Hodarchaeum class).

The class is delineated based on 209 concatenated AsCOGs and 16S rRNA gene phylogeny. The description is the same as that of its sole and type order *Candidatus* Hodarchaeales.

#### *Candidatus* Hodarchaeota

(Hod.ar.chae.o’ta. N.L. neut. n. Hodarchaeum a (Candidatus) type genus of the class of the phylum; -ota ending to denote the phylum; N.L neut. pl. n. Hodarchaeota the Hodarchaeum phylum)

The phylum is delineated based on 209 concatenated AsCOGs and 16S rRNA gene phylogeny. The description is the same as that of its sole and type class *Candidatus* Hodarchaeia.

#### *Candidatus* Kariarchaeaceae

(Ka.ri.ar.chae.a.ce’ae. N.L. neut. n. Kariarchaeum a (Candidatus) type genus of the family; -aceae ending to denote the family; N.L. fem. pl. n. Kariarchaeaceae the Kariarchaeum family).

The family is delineated based on 209 concatenated AsCOGs and 16S rRNA gene phylogeny. The description is the same as that of its sole genus and species. Type genus is *Candidatus* Kariarchaeum.

#### *Candidatus* Kariarchaeales

(Ka.ri.ar.chae.a’les. N.L. neut. n. Kariarchaeum a (Candidatus) type genus of the order; -ales ending to denote the order; N.L fem. pl. n. Kariarchaeales the Kariarchaeum order).

The order is delineated based on 209 concatenated AsCOGs and 16S rRNA gene phylogeny. The description is the same as that of its sole genus and species. Type genus is *Candidatus* Kariarchaeum.

#### *Candidatus* Kariarchaeia

(Ka.ri.ar.chae’i.a. N.L. neut. n. Kariarchaeum a (Candidatus) type genus of the order of the class; -ia ending to denote the class; N.L fem. pl. n. Kariarchaeia the Kariarchaeum class).

The class is delineated based on 209 concatenated AsCOGs and 16S rRNA gene phylogeny. The description is the same as that of its sole and type order *Candidatus* Kariarchaeales.

#### *Candidatus* Kariarchaeota

(Ka.ri.ar.chae.o’ta. N.L. neut. n. Kariarchaeum a (Candidatus) type genus of the class of the phylum; -ota ending to denote the phylum; N.L neut. pl. n. Kariarchaeota the Kariarchaeum phylum)

The phylum is delineated based on 209 concatenated AsCOGs and 16S rRNA gene phylogeny. The description is the same as that of its sole and type class *Candidatus* Kariarchaeia.

#### *Candidatus* Borrarchaeaceae

(Borr.ar.chae.a.ce’ae. N.L. neut. n. Borrarchaeum a (Candidatus) type genus of the family; -aceae ending to denote the family; N.L. fem. pl. n. Borrarchaeaceae the Borrarchaeum family).

The family is delineated based on 209 concatenated AsCOGs phylogeny. The description is the same as that of its sole genus and species. Type genus is *Candidatus* Borrarchaeum.

#### *Candidatus* Borrarchaeales

(Borr.ar.chae.a’les. N.L. neut. n. Borrarchaeum a (Candidatus) type genus of the order; -ales ending to denote the order; N.L fem. pl. n. Borrarchaeales the Borrarchaeum order).

The order is delineated based on 209 concatenated AsCOGs phylogeny. The description is the same as that of its sole genus and species. Type genus is *Candidatus* Borrarchaeum.

#### *Candidatus* Borrarchaeia

(Borr.ar.chae’i.a. N.L. neut. n. Borrarchaeum a (Candidatus) type genus of the order of the class; -ia ending to denote the class; N.L fem. pl. n. Borrarchaeia the Borrarchaeum class).

The class is delineated based on 209 concatenated AsCOGs phylogeny. The description is the same as that of its sole and type order *Candidatus* Borrarchaeales.

#### *Candidatus* Borrarchaeota

(Borr.ar.chae.o’ta. N.L. neut. n. Borrarchaeum a (Candidatus) type genus of the class of the phylum; -ota ending to denote the phylum; N.L neut. pl. n. Borrarchaeota the Borrarchaeum phylum)

The phylum is delineated based on 209 concatenated AsCOGs phylogeny. The description is the same as that of its sole and type class *Candidatus* Borrarchaeia.

#### *Candidatus* Baldrarchaeaceae

(Bal.dr.ar.chae.a.ce’ae. N.L. neut. n. Baldrarchaeum a (Candidatus) type genus of the family; -aceae ending to denote the family; N.L. fem. pl. n. Baldrarchaeaceae the Baldrarchaeum family).

The family is delineated based on 209 concatenated AsCOGs and 16S rRNA gene phylogeny. The description is the same as that of its sole genus and species. Type genus is *Candidatus* Baldrarchaeum.

#### *Candidatus* Baldrarchaeales

(Bal.dr.ar.chae.a’les. N.L. neut. n. Bladrarchaeum a (Candidatus) type genus of the order; -ales ending to denote the order; N.L fem. pl. n. Baldrarchaeales the Baldrarchaeum order).

The order is delineated based on 209 concatenated AsCOGs and 16S rRNA gene phylogeny. The description is the same as that of its sole genus and species. Type genus is *Candidatus* Baldrarchaeum.

#### *Candidatus* Baldrarchaeia

(Bal.dr.ar.chae’i.a. N.L. neut. n. Baldrarchaeum a (Candidatus) type genus of the order of the class; -ia ending to denote the class; N.L fem. pl. n. Baldrarchaeia the Baldrarchaeum class).

The class is delineated based on 209 concatenated AsCOGs and 16S rRNA gene phylogeny. The description is the same as that of its sole and type order *Candidatus* Baldrarchaeales.

#### *Candidatus* Baldrarchaeota

(Bal.dr.ar.chae.o’ta. N.L. neut. n. Baldrarchaeum a (Candidatus) type genus of the class of the phylum; -ota ending to denote the phylum; N.L neut. pl. n. Baldrarchaeota the Baldrarchaeum phylum)

The phylum is delineated based on 209 concatenated AsCOGs and 16S rRNA gene phylogeny. The description is the same as that of its sole and type class *Candidatus* Baldrarchaeia.

#### *Candidatus* Hermodarchaeaceae

(Her.mod.ar.chae.a.ce’ae. N.L. neut. n. Hermodarchaeum a (Candidatus) type genus of the family; -aceae ending to denote the family; N.L. fem. pl. n. Hermodarchaeaceae the Hermodarchaeum family).

The family is delineated based on 209 concatenated AsCOGs and 16S rRNA gene phylogeny. The description is the same as that of its sole genus and species. Type genus is *Candidatus* Hermodarchaeum.

#### *Candidatus* Hermodarchaeales

(Her.mod.ar.chae.a’les. N.L. neut. n. Hermodarchaeum a (Candidatus) type genus of the order; -ales ending to denote the order; N.L fem. pl. n. Hermodarchaeales the Hermodarchaeum order).

The order is delineated based on 209 concatenated AsCOGs and 16S rRNA gene phylogeny. The description is the same as that of its sole genus and species. Type genus is *Candidatus* Hermodarchaeum.

#### *Candidatus* Hermodarchaeia

(Her.mod.ar.chae’i.a. N.L. neut. n. Hermodarchaeum a (Candidatus) type genus of the order of the class; -ia ending to denote the class; N.L fem. pl. n. Hermodarchaeia the Hermodarchaeum class).

The class is delineated based on 209 concatenated AsCOGs and 16S rRNA gene phylogeny. The description is the same as that of its sole and type order *Candidatus* Hermodarchaeales.

#### *Candidatus* Hermodarchaeota

(Her.mod.ar.chae.o’ta. N.L. neut. n. Hermodarchaeum a (Candidatus) type genus of the class of the phylum; -ota ending to denote the phylum; N.L neut. pl. n. Hermodarchaeota the Hermodarchaeum phylum)

The phylum is delineated based on 209 concatenated AsCOGs and 16S rRNA gene phylogeny. The description is the same as that of its sole and type class *Candidatus* Hermodarchaeia.

### Materials and Methods

#### Sampling collections and DNA sequencing

YT samples were obtained from the Rongcheng Nation Swan Nature Reserve (Rongcheng, China) in November 2018. The sediment cores were collected using columnar samplers at depth intervals of 0–2, 21–26, and 36–41 cm at a seagrass meadow and a non-seagrass–covered site nearby. After collection, bulk sediments were immediately sealed in plastic bags, placed in a pre-cooled icebox, and transported to laboratory within 4 hours. For each sample, DNA was extracted from 10 g sediment using PowerSoil DNA Isolation kit (QIAGEN, Germany), according to the manufacturer’s protocol. Following extraction, nucleic acids were sequenced using Illumina HiSeq2500 (Illumina, USA) PE150 by Novogene (Nanjing, China).

MP5 samples were obtained from Mai Po Nature Reserve (Hong Kong, China) in September 2014. Three subsurface sediment samples were collected from a site covering with mangrove forest at depth intervals of 0-2, 10-15 and 20-25 cm. Two subsurface sediment samples were taken at an intertidal mudflat with depths of 0-5 and 13-16 cm. Samples were transported back to laboratory as described for YT metagenomes. DNA was extracted from 5g wet sediment per sample using the PowerSoil DNA Isolation Kit (MO BIO, USA) following the manufacturer’s protocol. Metagenomic sequencing data were generated using Illumina HiSeq2500 (Illumina, USA) PE150 by Novogene (Tianjin, China).

FT samples were taken from Futian Nature Reserve (Shenzhen, Guangdong, China) in April 2017. Sediment samples were collected as described for YT samples at depth intervals of 0-2, 6-8, 12-14, 20-22, and 28-30 cm. DNA was extracted from 5g wet sediment per sample using DNeasy PowerSoil kit (Qiagen, Germany) as per manufacturer’s instructions. Nucleic acids were sequenced using Illumina HiSeq2000 (Illumina, USA) PE150 by Novogene (Tianjin, China).

The surface sediment sample of the CJE metagenome was collected from Changjiang estuary during a cruise in August 2016. The sample was grabbed from the water bed, sealed immediately in a 50 ml tubes and stored in liquid nitrogen onboard. After transportation to laboratory, 10g wet sediments were used for DNA extraction as per manufacturer’s protocol. Nucleic acids were sequenced using Illumina HiSeq2000 (Illumina, USA) PE150 by Novogene (Tianjin, China).

An oil sand sample was collected from Shengli Oilfield (Shandong, China) into bottles and they were transported to laboratory where they were stored at 4 °C. The sample was used as inoculum to perform enrichment with anaerobic medium in vials as described elsewhere (*1*). After 253d of enrichment, the genomic DNA was extracted as described elsewhere (*2*). Nucleic acids were sequenced Illumina HiSeq2000 (Illumina, USA) PE150 by Novogene (Tianjin, China).

The seawater samples of Yap metagenomes were collected at Yap trench region by CTD SBE911plus (Sea-Bird Electronics, USA) during the 37^th^ Dayang cruise in 2016. 8L of seawater per sample was filtered through a 0.22 μm-mesh membrane filter immediately after recovery onboard. The membrane was then cut into ~0.2 cm^2^ pieces with a flame-sterilized scissors and added to a PowerBead Tube (MO BIO, USA) and the subsequent steps were implemented according to the manufacturer’s protocol to extract DNA. The DNA per sample was amplified in five separative reactions using REPLI-g Single Cell Kits (Qiagen, Germany) following the manufacturer’s protocol, given to the challenging nature of sample retrieval and DNA recovery. The products were pooled together and purified using QIAamp DNA Mini Kit (Qiagen, Germany) according to the manufacturer’s recommendations. Parallel blank controls were set for sampling, DNA extraction and ampliation with 0.22 μm-mesh membrane filtering Milli-Q water (18.2 MΩ; Millipore, USA). Nucleic acids were sequenced using HiSeq X Ten (Illumina, USA) PE150.

#### Metagenomic assembly, binning and gene calling

##### FT, MP5 and YT metagenomes

These three sets of metagenomes were assembled and binned using the same method. Raw shotgun metagenomic sequencing reads were trimmed with “read_qc” module from metaWRAP (v.1.1) (*3*). All clean reads from the same set were pooled together prior to *de novo* assemble to one co-assembly. Clean reads were sent out to MEGAHIT (v1.1.2) with flag “--presets meta-large” for co-assembling job (*4*). Sequencing coverage was determined for each assembled scaffold by mapping reads from each sample to the co-assembly using Bowtie2 (*5*). The binning analysis was carried out 8 times with 8 different combinations of specificity and sensitivity parameters using MetaBAT2(52) (“-- maxP 60 or 95” AND “--minS 60 or 95” AND “--maxEdges 200 or 500”) on the assembly with a minimum length of 2000 bp (*6*). DAS Tool (v1.0) was used as a dereplication and aggregation strategy on those eight binning results to construct accurate bins (*7*). Manual curation was used for reducing the genome contamination based on differential coverages, GC contents, and the presence of duplicate genes.

The depth coverage and N50 statistics of 38 Asgard MAGs recovered from YT metagenomes range from 7.72 to 298.86 (median: 21.77) and from 6658 to 381755 bps (median: 26337.5 bps) respectively; for 13 Asgard MAGs recovered from FT metagenomes, the values range from 6.98 to 33.22 (median 16.89) and from 3889 to 8957 bps (median: 5000 bps), respectively; and for 11 Asgard MAGs recovered from MP5 metagenomes, the values range from 7.06 to 54.42 (median: 10) and from 3898 to 19362 bps (median: 7581 bps) respectively. (**Supplementary Figure 2**).

##### CJE metagenome

Raw metagenomic shotgun sequencing reads were trimmed using Sickle (https://github.com/najoshi/sickle) with default settings. The trimmed reads were de novo assembled using IDBA-UD (v 1.1.1) with the parameters: “-mink 65, -maxk 145, -step 10” (*8*). Sequencing coverage was determined as described above. The binning analysis were performed with MetaBAT2 12 times, with 12 combinations of specificity and sensitivity parameters (--m 1500, 2000, or 2500” AND “--maxP 85 or 90” AND “--minS 80, 85 and 90”) for further refinement (*6*). All binning results were merged and refined using DAS Tool (v1.0) (*7*).

2 Asgard MAGs recovered from CJE metagenome have a depth coverage of 19.16 and 20.56, and a N50 statistics of 8740 and 5246 bps (**Supplementary Figure 2**).

##### J65 metagenome

Raw metagenomic shotgun sequencing reads were trimmed with Trimmomatic (v0.38) (*9*). The clean reads were then fed to SPAdes (v 3.12.0) for *de novo* assembly with the parameters: “-- meta -k 21, 33, 55, 77” (*10*). Sequencing coverage was determined using BBMap (v 38.24) toolkit with the parameters: “bbmap.sh minid=0.99” (https://github.com/BioInfoTools/BBMap). MetaBAT2 (v 2.12.1) was used to perform binning analysis with the parameter: “-m 2000” (*6*).

One Asgard MAG recovered from J65 metagenome had a depth coverage of 14.52 and a N50 statistics of 10460 bps (**Supplementary Figure 2**).

##### Yap metagenome

For each Yap metagenome, raw metagenomic shotgun sequencing reads were trimmed with Trimmomatic (v0.38) (*9*). Assembly and binning analysis were performed as described for CJE metagenome for each Yap metagenome.

The depth coverage and N50 statistics of 24 Asgard MAGs recovered from Yap metagenomes range from 7.78 to 82.33 (median: 13.91) and from 5097 to 889102 bps (median: 18155.5 bps) respectively. (**Supplementary Figure 2**)

A total of 89 Asgard MAGs were reconstructed in this work. Additional 95 Asgard MAGs were downloaded from public databases (e.g. NCBI FTP site). For all 184 genomes, a uniform gene calling protocol was applied. Specifically, the completeness, contamination, and strain heterogeneity of the genomes were estimated by using CheckM (v.1.0.12) (*11*) and DAS Tool under the taxonomic scope of domain (i.e., Bacteria and Archaea). Protein-coding genes were predicted using Prodigal (v 2.6.3) (*12*) embedded in Prokka (v 1.13) (*13*). Transfer RNAs (tRNAs) were identified with tRNAscan-SE (v1.23) using the archaeal tRNA model (*14*). After quality screening, further analysis focused on 162 high quality Asgard MAGs.

#### Genome set

The Asgard archaea genome set analyzed in this work consisted of 161 Asgard MAGs and one complete Asgard genome (**Supplementary Table 1**). For comparison, selected representative genomes of archaea (296), bacteria (100) and eukaryotes (76) were downloaded from Refseq and Genbank using the NCBI FTP site (ftp://ftp.ncbi.nlm.nih.gov/genomes/all/).

#### Average amino-acid identity

The average amino-acid identity (AAI) across TACK archaeal reference genomes and the 184 Asgard genomes was calculated using compareM (v0.0.23) with the “aai_wf” at default settings (https://github.com/dparks1134/CompareM).

#### asCOGs construction

Initial clustering of 250,634 proteins encoded in 76 Asgard MAGs was performed using two approaches: first, footprints of arCOG profiles were obtained by running PSI-BLAST (*15*), initiated with arCOG alignments, against the set of predicted Asgard proteins. The footprint sequences were extracted and clustered according to the arCOG best hit. The remaining protein sequences (both full-length proteins and the sequence fragments outside of the footprints, if longer than 60 aa) were clustered using MMseqs2 (*16*), with similarity threshold of 0.5. Sequences within clusters were aligned using MUSCLE (*17*); the resulting alignments were passed through several rounds of merging and splitting. The merging phase involved comparing alignments to each other using HHSEARCH (*18*), finding full-length cluster-to-cluster matches, merging the sequence sets and re-aligning the new clusters. The splitting phase consisted of the construction of an approximate phylogenetic tree of the sequences using FastTree (*19*) (gamma-distributed site rates, WAG evolutionary model) with balanced mid-point tree rooting, identification of subtrees maximizing the fraction of species (MAGs) representation and minimizing the number of paralogs, and pruning such subtrees as separate clusters of putative orthologs. Clusters, derived from arCOGs, were prohibited from merging across distinct arCOGs to prevent distant paralogs from forming mixed clusters.

#### Phylogenetic analysis

##### 16S rRNA gene phylogenetic analysis

16S rRNA gene sequences were identified in 73 genomes of Asgard archaea (26 generated in this study, 47 from public database) using Barrnap (v 0.9) with the “-- kingdom arc” option (https://github.com/tseemann/barrnap). These sequences were combined with 46 published 16S rRNA sequences of Asgard archaea, to assess the novelty of the sequences obtained in this work. The novelty of 16S rRNA gene sequences was measured in terms of their sequence identity to previously identified Asgard archaeal 16S rRNA gene and phylogenetic relationships. Specifically, the pairwise sequence identity of two Asgard archaeal 16S rRNA gene sequences (>1300bp) was obtained by first globally aligning the sequences with “Stretcher” in EMBOSS package and then calculating the percent identity excluding gaps (*20*). The Asgard archaeal 16S rRNA gene sequences were aligned with 311 reference sequences from Euryarchaeota (n=231), DPANN (n=22), Korarchaeota (n=1), Crenarchaeota (n=41), Bathyarchaeota (n=2), Thaumarchaeota (n=13) and Aigarchaeota (n=1) using mafft-linsi (v7.471) (*21*) and trimmed with BMGE (v1.12) (settings: -m DNAPAM250:4, -g 0.5) (*22*).The alignment was used for phylogenetic inference with IQ-Tree (v2.0.6) based on SYM+R8 (selected by ModelFinder) to generate a maximum-likelihood tree (*23*).

##### Asgard phylogeny

A set of asCOGs that were considered most suitable as phylogenetic markers for Asgard archaea was selected using the preliminary classification of the 76 genomes in AsCOGs into previously described lineages: Loki, Thor, Odin, Hel and Heimdall. The following criteria were adopted: the asCOG have to be i) present in at least half of the genomes in all lineages, ii) present in at least 75% among the 76 genomes, iii) the mean number of paralogs per genome not to exceed 1.25. For the 209 asCOGs matching these criteria, the corresponding protein sequences were obtained from the extended set of 162 MAGs and aligned using MUSCLE (*17*); the ‘index’ paralog to include in the phylogeny was selected for each MAG based on the similarity to the alignment consensus. Alignments were trimmed to exclude columns containing more than 2/3 gap characters and with homogeneity below 0.05 (homogeneity is calculated from the score of the consensus amino acid against the alignment column, compared to the score of the perfect match (*24*) and concatenated, resulting in an alignment of 50,706 characters from 162 sequences (one sequence per MAG). Phylogeny was reconstructed using FastTree (gamma-distributed site rates, WAG evolutionary model) (*19*) and IQ-Tree (LG+F+R10 model, selected by ModelFinder) (*23*), producing very similar tree topologies.

##### Tree of life

To elucidate the relationships between the Asgard archaea and other major clades of archaea, bacteria and eukaryotes, 30 families of conserved proteins were selected that appear to have evolved mostly vertically (25, 26). The set of 162 MAGs of Asgard archaea was supplemented with 66 TACK archaea and 220 non-TACK archaea (the former, having been described as the closest archaeal relatives of the Asgard archaea (*27*), were sampled more densely), 98 bacteria and 72 eukaryotes. Prokaryotic genomes were sampled from the set of completely sequenced genomes to represent the maximum diversity within their respective clades (briefly, all proteins, encoded by genomes within a groups were clustered at 75% identity level; distances between genomes were estimated from the number of shared proteins within these clusters; UPGMA trees were reconstructed from the distances, and genome sets maximizing the total tree branch length were selected to represent the groups). The set of eukaryotic genomes was manually selected to represent the maximum possible variety of eukaryotic taxa. Genomes, in which more than four markers were missing, were excluded from the bacterial and eukaryotic sets. When multiple paralogs of a marker were present in a genome, preliminary phylogenetic trees were constructed from protein sequence alignments, and paralogs with the shortest branches were selected to represent the corresponding genomes in the set. Sequences were aligned using MUSCLE (*17*), and alignment columns, containing more than 2/3 gap characters or with alignment column homogeneity below 0.05 were removed (*24*). The resulting concatenated alignments of the 30 markers consisted of 7411 sites. Phylogeny was reconstructed using FastTree (gamma-distributed site rates, WAG evolutionary model) (*19*) and IQ-Tree (*23*) with three models: LG+R10, selected by IQ-tree ModelFinder as the best fit, GTR20+F+R10 (following the suggestion of Williams et al. (*28*) to use GTR, we let IQ-tree to select the best version of the GTR model), and LG+C20+G4+F (again, the mixture model was used following the suggestion of Williams et al. (*28*); we were unable to use the higher-specified C60 model due to memory limitations of our hardware and used the C20 model instead).

#### Ordination of asCOG phyletic patterns using Classical Multidimensional Scaling

Binary asCOG presence-absence patterns were compared between pairs of Asgard MAGs using the following procedure: first, similarity between asCOG sets {***A***} and {***B***} was calculated as 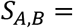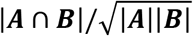 (the number of shared AsCOGs normalized by the geometric mean of the number of AsCOGs in the two MAGs); then, the distance between the patterns was calculated as *d*_*A,B*_ = −ln(*S*_*A,B*_). The 162×162 distance matrix was embedded into a 2-dimensional space using Classical Multidimensional Scaling analysis implemented as the *cmdscale* function in R. The projection retained 89% of the original datapoint inertia.

#### Identification and analysis of ESPs

To identify eukaryotic signature proteins (ESPs), several strategies were employed. First, ESPs reported by Zaremba-Niedzwiedzka et al. (*29*) were mapped to asCOGs using PSI-BLAST (*15*). These asCOGs were additionally examined case by case using HHpred (*30*) with representative of the respective asCOG or the respective asCOG alignment used as the query. Second, all asCOGs were mapped to CDD profiles (*31*) using PSI-BLAST, hits to eukaryote-specific domains were selected. Most of the putative ESP asCOGs identified in this search, and all with E-value >1e-10, were additionally examined using HHpred, with a representative of the respective asCOG or the respective asCOG alignment as the query. Third, we analyzed frequently occurring asCOGs (present in at least 50% of Asgard genomes and in at least 30% of Heimdall genomes) that were not annotated automatically with the above two approaches using HHpred with a representative of the respective asCOG or the respective asCOG alignment as the query. Fourth, most of the putative ESP asCOGs detected with these approaches were used as queries for a PSI-BLAST search that was run for three iterations (with inclusion threshold E-value=0.0001) against an Asgard only protein sequence database. Additional unannotated asCOGs with similarity to the (putative) ESPs identified in this search were further examined using HHpred. Fifth, the genomic neighborhoods or all ESPs were examined, and proteins encoded by unannotated neighbor genes were analyzed using HHpred server.

#### Metabolic pathway reconstruction

The patterns of gene presence-absence in the asCOGs were used to reconstruct the metabolic pathways of Asgard archaea (Supplementary Table 8). The asCOGs were linked to the KEGG database and to the list of predicted metabolic enzymes of Asgard archaea reported by Spang et al. (*32*) (Supplementary Table 8). The classification of [NiFe] hydrogenases was performed by comparing the Asgard proteins of cog.001539, cog.002254, cog.010021, cog.011939 and cog.012499 to HydDB (*33*). For phylogenetic analysis, the reference sequences of group 1, 3 and 4 [NiFe] hydrogenases were retrieved from HydDB. The sequences were filtered using cd-hit with a sequence identity cut-off of 90% prior to adding orthologous genes of cog.001539, cog.002254, cog.010021, cog.011939 and cog.012499 of Asgard archaea. All sequences for group 1, group 3 and group 4 [NiFe] hydrogenases were aligned using mafft-LINSI (*21*) and trimmed with BMGE (-m BLOSUM30 -h 0.6) (*22*). Maximum-likelihood phylogenetic analyses were performed using IQ-tree (*23*) with the best-fit model (group 1:LG + C60 + R + F, group 3: LG + C60 + R +F and group 4 LG + C50 + R + F), respectively, according to Bayesian information criterion (BIC). Support values were estimated using the SH-like approximate-likelihood ratio test and ultrafast bootstraps, respectively. We adapted a relaxed common denominator approach to determine the presence of certain pathway in one Asgard phyla (*32*), and combined with maximum parsimony principle (*34*) to infer the metabolisms of major ancestral forms.

## Data availability

Asgard archaea genomes generated in this study are available in eLMSG (an eLibrary of Microbial Systematics and Genomics, https://www.biosino.org/elmsg/index) under accession numbers from LMSG_G000000521.1 to LMSG_G000000609.1 and NCBI database under the project XXXXX.

Additional Data files are available at ftp://ftp.ncbi.nih.gov/pub/wolf/_suppl/asgard20/

Additional data file 1 (Additional_data_file_1.tgz): Complete asCOG data archive

Additional data file 2 (Additional_data_file_2.tgz): Phylogenetic trees and alignments archive

## Supplementary Figures

**Supplementary Figure 1.**
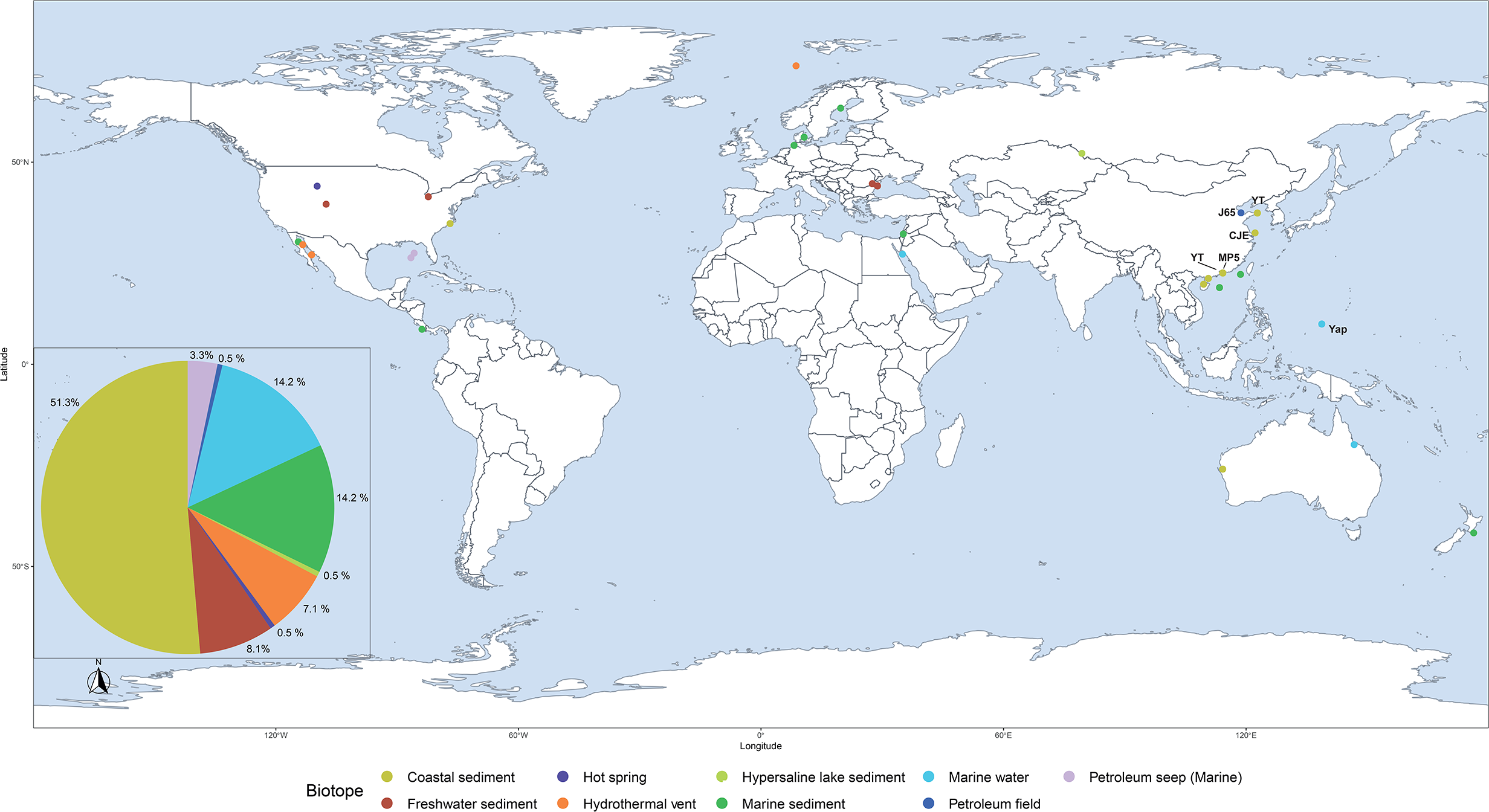
Global distribution of Asgard genomes analyzed in the study. The pie chart shows the proportion of Asgard genomes found in a biotope. Bold letters in the map show the sampling locations.

**Supplementary Figure 2.**
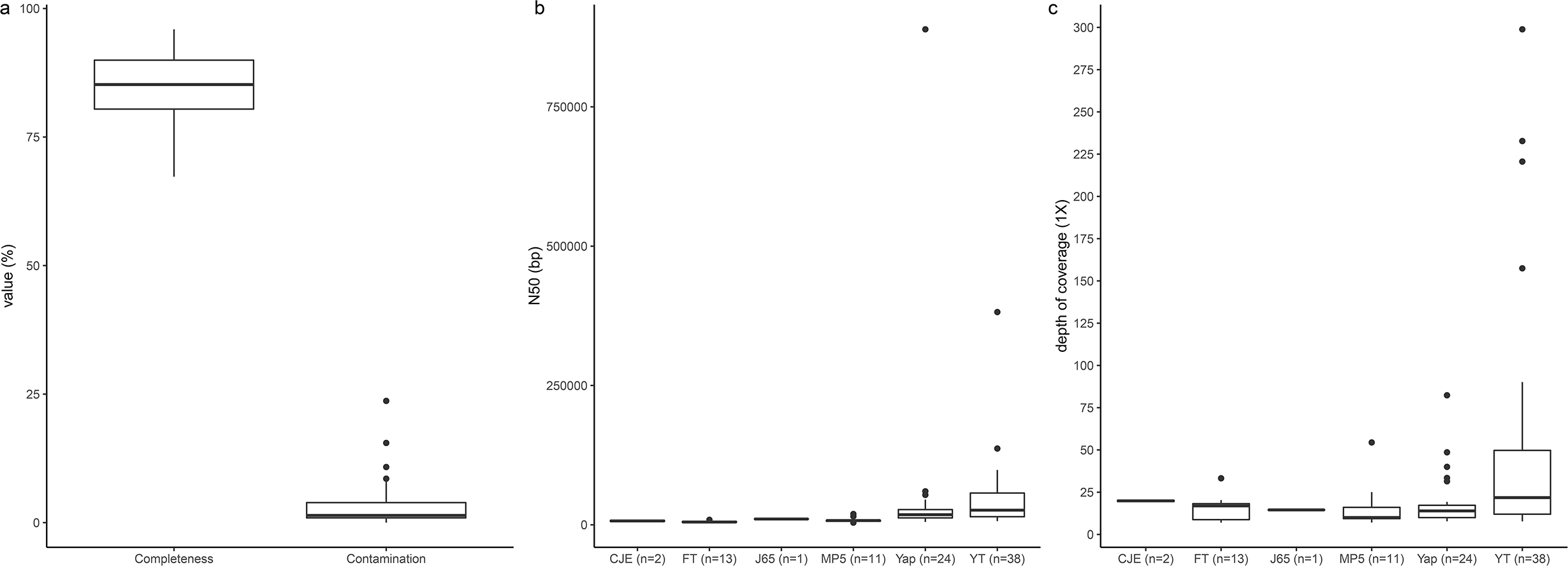
a. Distribution of completeness and contamination for 76 Asgard MAGs assessed by CheckM (v 1.0.12). Distribution of depth coverage (b) and N50 statistics (c) for Asgard MAGs reconstructed in this study. The numbers in parentheses indicate the number of Asgard genomes recovered from a given sampling location. The data for this plot can be found in **Supplementary Table 1.**

**Supplementary Figure 3.**
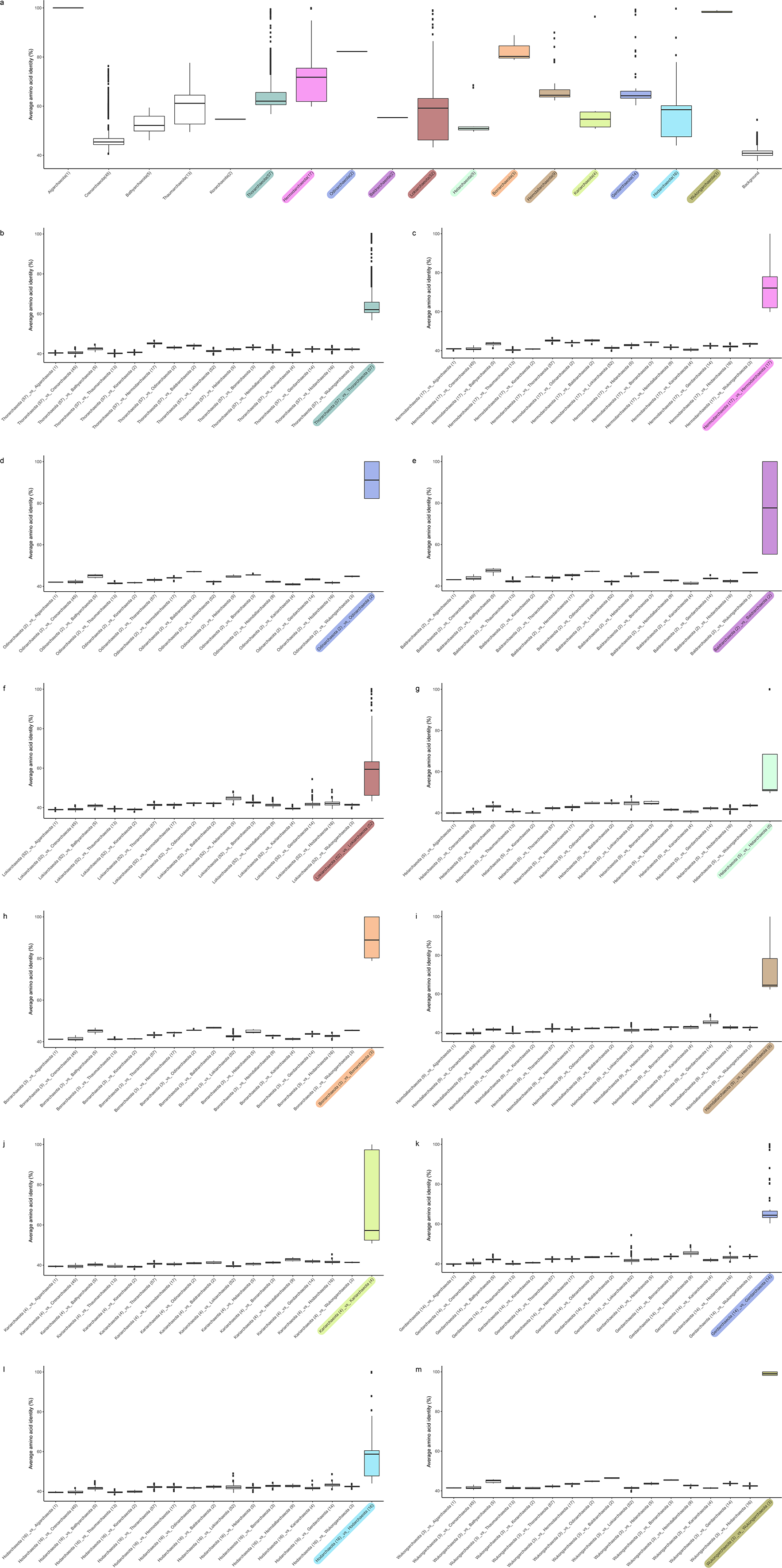
Comparison of the mean amino-acid identity (AAI) of Asgard and TACK superphyla. a. Shared AAI across Asgard and TACK lineages. Background: comparing all Asgard and TACK lineages included in the analyses but excluding archaea belonging to the same lineages and the six phyla proposed in the current work to investigate the distribution of AAI that defines a phylum. The AAI comparison of (b) Thorarchaeota, (c) Hermodarchaeota (d) Odinarchaeota, (e) Baldrarchaeota, (f) Lokiarchaeota, (g) Helarchaeota, (h) Borrarchaeota, (i) Heimdallarchaeota, (j) Kariarchaeota, (k) Gerdarchaeota, (l) Hodarchaeota and (m) Wukongarchaeota to other Asgad and TACK lineages. The lower and upper hinges of the boxplot correspond to the first and third quartiles. Data beyond the whiskers are shown as individual data points. Number in the parenthesis indicates the number of genomes in the lineages. Data for this plot is included in **Supplementary Table 2.**

**Supplementary Figure 4.**
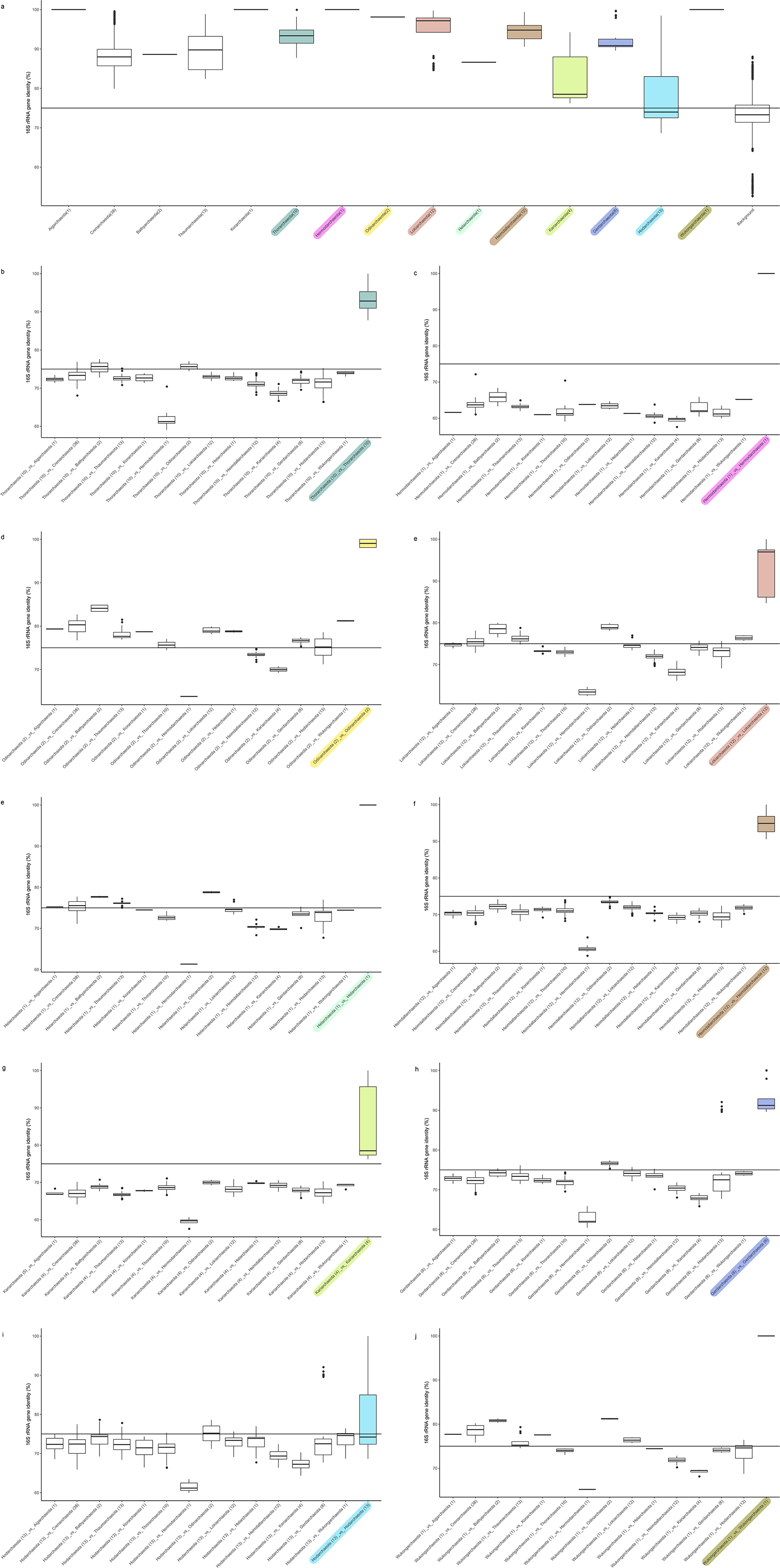
Comparison of the 16S rRNA gene sequence (>1300 bp) identity of Asgard and TACK lineages. a. Shared 16S rRNA gene sequence identity across Asgard and TACK lineages. Background:comparing all Asgard and TACK lineages included in the analyses but excluding archaea belonging to the same lineages and the six phyla proposed in the current study to investigate the distribution of 16S rRNA gene identity that defines a phylum. 16S rRNA gene identity comparison of (b) Thorarchaeota, (c) Hermodarchaeota (d) Odinarchaeota, (e) Lokiarchaeota, (f) Helarchaeota, (g) Heimdallarchaeota, (h) Kariarchaeota, (i) Gerdarchaeota, (j) Hodarchaeota and (k) Wukongarchaeota to other Asgard and TACK lineages. The lower and upper hinges of the boxplot correspond to the first and third quartiles. Data beyond the whiskers are shown as individual data points. Number in the parenthesis indicates the number of 16S rRNA gene sequences compared in the lineages. Line represents a 16S rRNA gene identity of 75%. Data for this plot could be found in **Supplementary Table 3.**

**Supplementary Figure 5.**
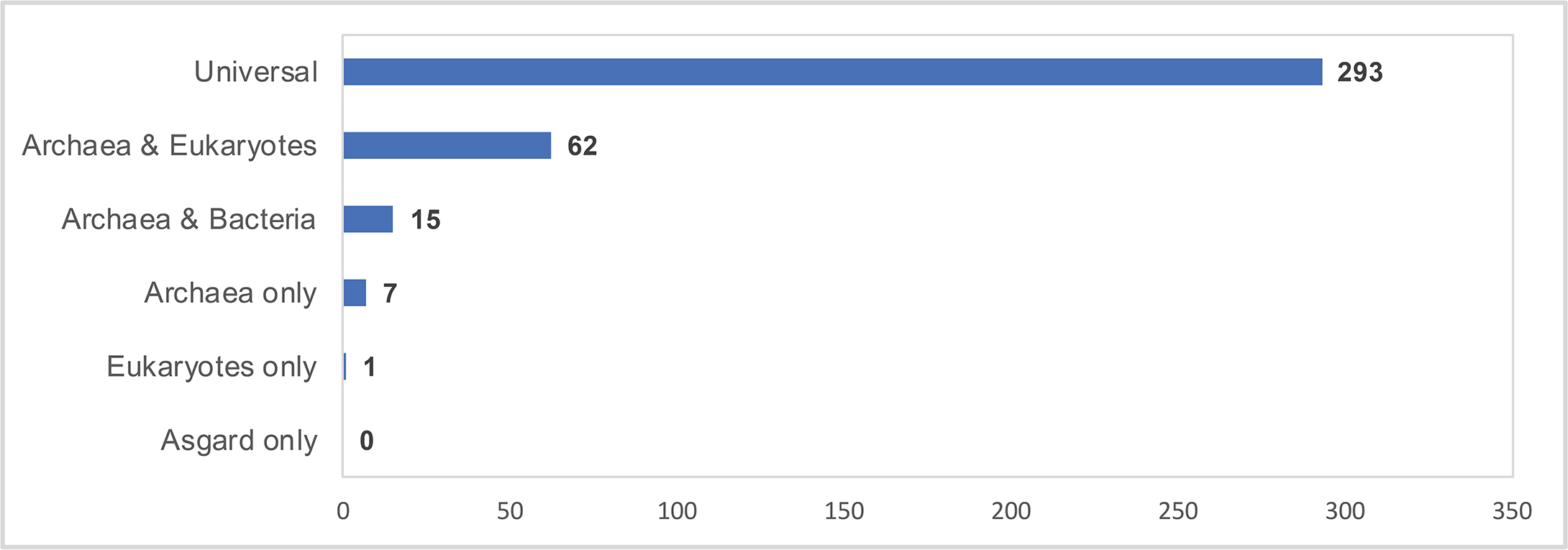
Presence-absence of orthologs of Asgard core genes in other archaea, bacteria and eukaryotes.

**Supplementary Figure 6.**
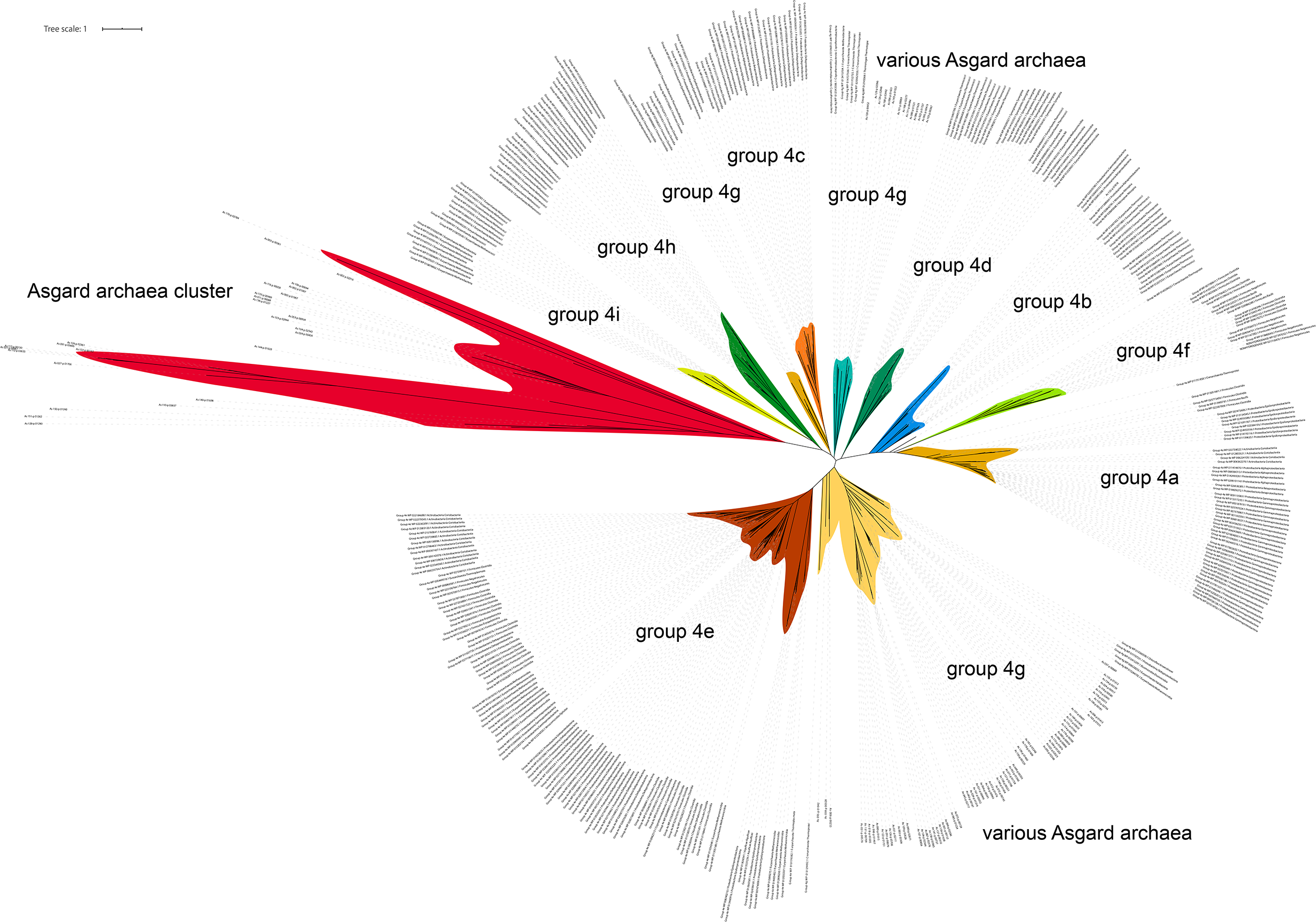
Phylogenetic analysis of group 4 [NiFe] hydrogenases in the Asgard archaea. The unrooted maximum-likelihood phylogenetic tree was built from an alignment of 425 sequences including 110 sequences of Asgard archaea, with 308 amino-acid positions.

**Supplementary Figure 7.**
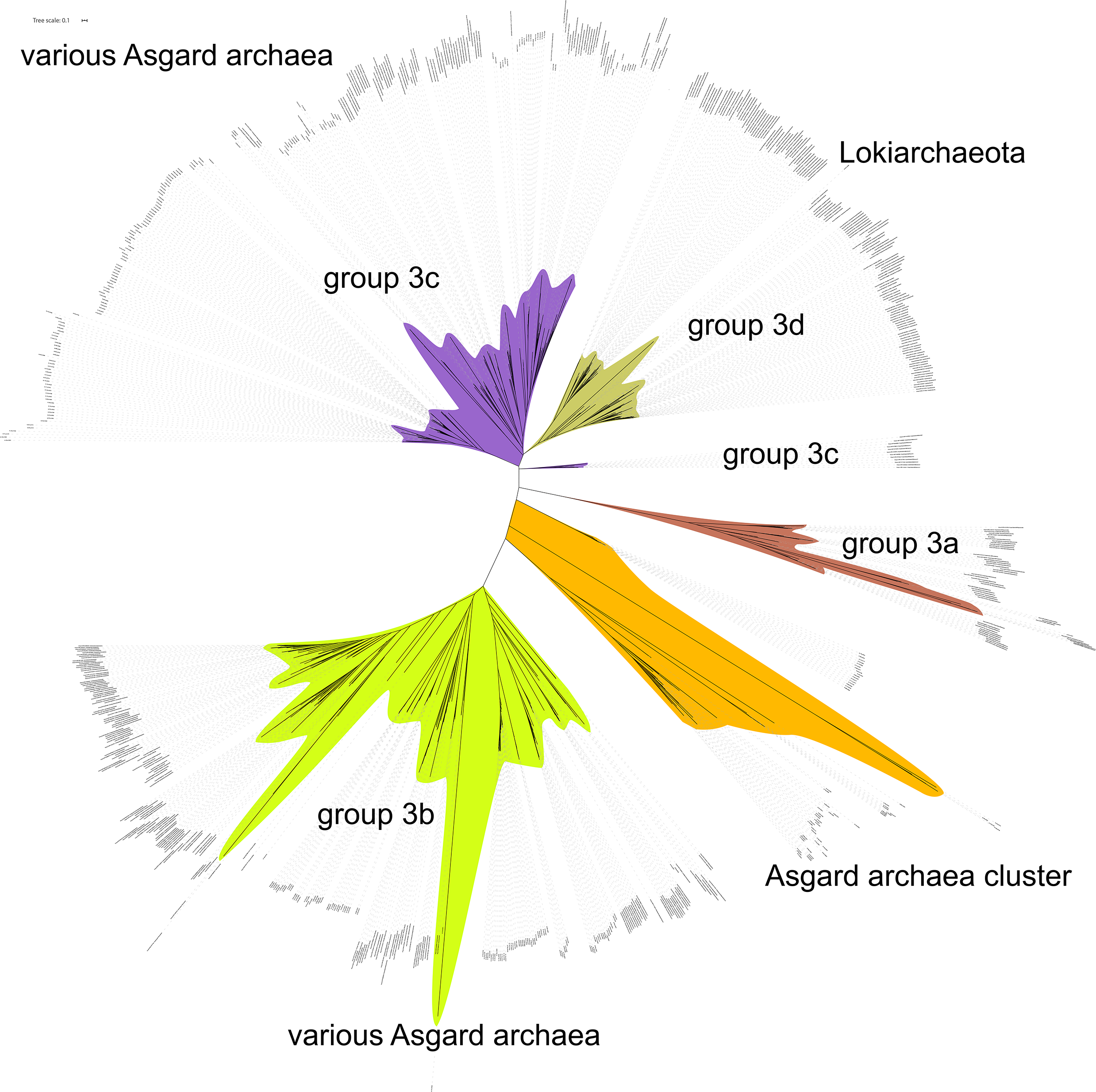
Phylogenetic analysis of group 3 [NiFe] hydrogenases in the Asgard archaea. The unrooted maximum-likelihood phylogenetic tree was built from an alignment of 813 sequences including 335 sequences of Asgard archaea, 514 amino-acid positions.

**Supplementary Figure 8.**
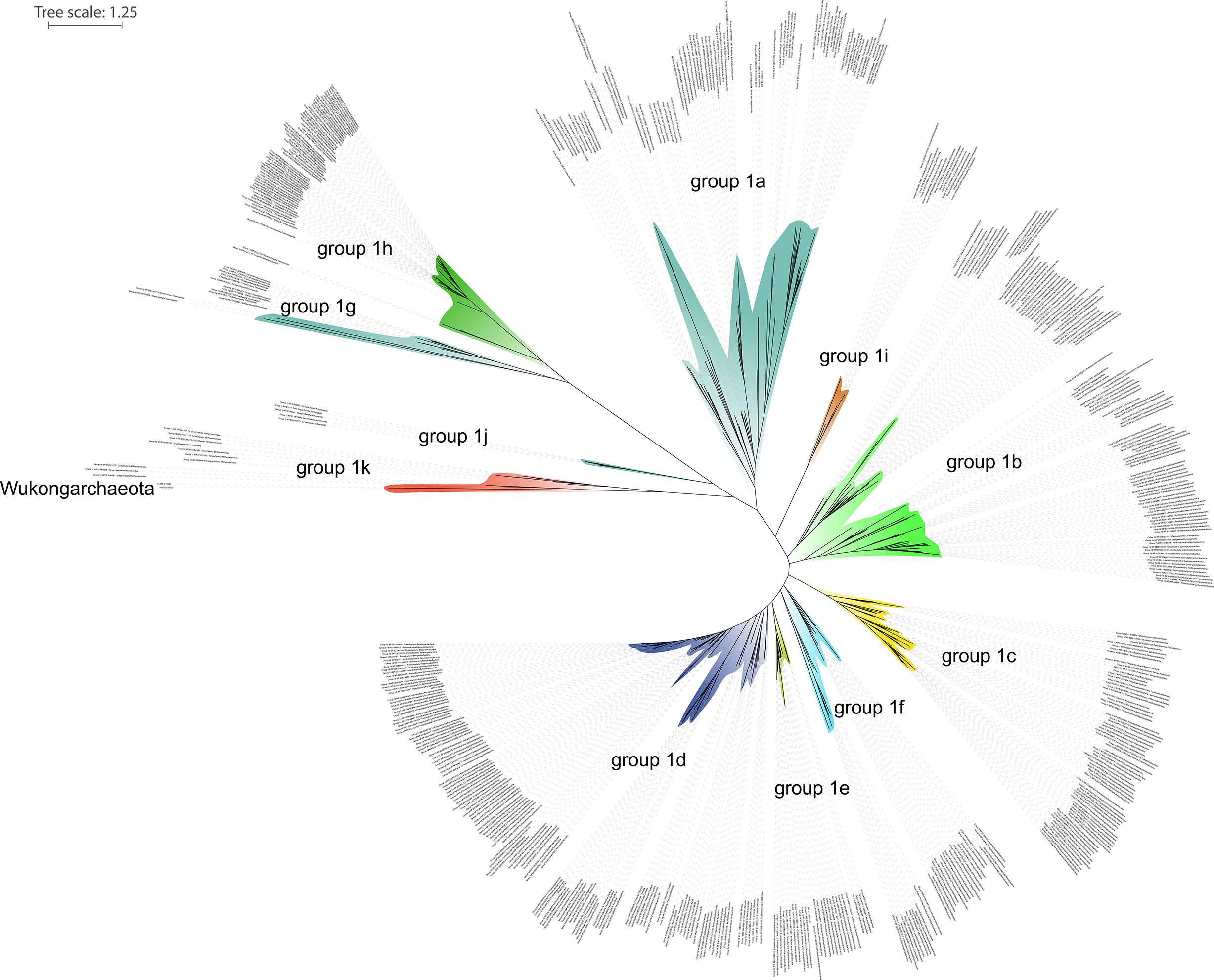
Phylogenetic analysis of group 1 [NiFe] hydrogenases in the Asgard archaea. The unrooted maximum-likelihood phylogenetic tree was built from an alignment of 541 sequences including 2 sequences of Wukongarchaeaota, with 745 amino-acid positions.

**Supplementary Figure 9.**
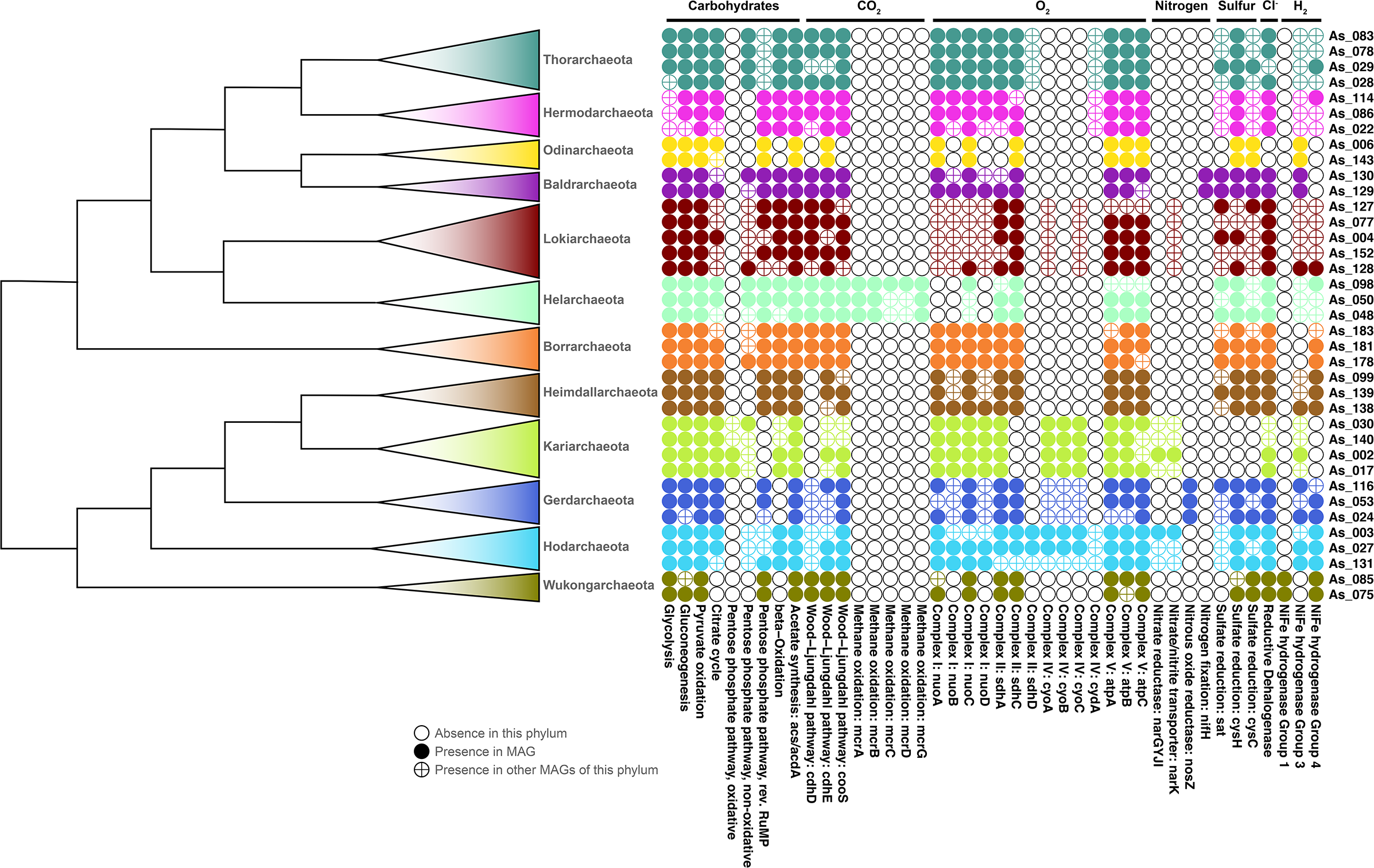
A schematic representation of the presence and absence of selected metabolic features in all (putative) phyla of Asgard archaea.

**Supplementary Figure 10.**
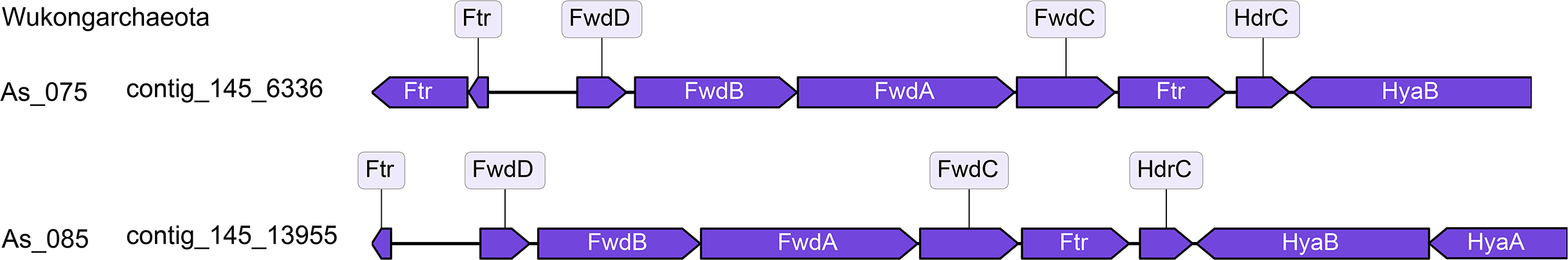
Gene structure of the contig encoding Group 1 [Ni,Fe]-hydrogenase in Wukongarchaeota. Abbreviations: Ftr, Formylmethanofuran:tetrahydromethanopterin formyltransferase; FwdD Formylmethanofuran dehydrogenase subunit D; FwdB, Formylmethanofuran dehydrogenase subunit B; FwdA, Formylmethanofuran dehydrogenase subunit A; FwdC, Formylmethanofuran dehydrogenase subunit C; HdrC, Heterodisulfide reductase, subunit C; HyaB, Ni,Fe-hydrogenase I large subunit; HyaA, Ni,Fe-hydrogenase I small subunit.

## Supplementary Tables

**Supplementary Table 1** Genome information, proposed taxonomy and isolation data

**Supplementary Table 2** Mean amino-acid identity values (in %) comparing 66 TACK genomes and 184 Asgard genomes (162 high quality and 22 low-quality)

**Supplementary Table 3** The 16S rRNA gene sequence identity (in %) comparing TACK lineages and Asgard lineages. The identity was calculated using sequences longer than 1300 bps

**Supplementary Table 4** Species and phyletic markers used for the tree of life reconstruction

**Supplementary Table 5** Data for phylogenetic trees: the trees in the Newick format and the underlying alignments

**Supplementary Table 6** The asCOGs annotation

**Supplementary Table 7** Eukaryotic signature proteins in Asgard archaea

**Supplementary Table 8** The presence-absence of metabolic enzymes in Asgard archaea.

